# Synthetic Biodegradable Void-forming Hydrogels for *In Vitro* 3D Culture of Functional Human Bone Cell Networks

**DOI:** 10.1101/2023.10.23.563580

**Authors:** Doris Zauchner, Monica Zippora Müller, Marion Horrer, Leana Bissig, Feihu Zhao, Sung Sik Lee, Ralph Müller, Xiao-Hua Qin

## Abstract

Generating 3D bone cell networks *in vitro* that accurately mimic the dynamic process of osteoblast embedding during early bone formation poses a significant challenge. Herein, we report a synthetic biodegradable macroporous hydrogel for efficient formation of 3D networks from human primary cells, analysis of cell-secreted extracellular matrix (ECM) and microfluidic integration. Using polymerization-induced phase separation, matrix metalloproteinase-sensitive polyethylene glycol hydrogels are formed with interconnected porosity in the presence of living cells. The pore size (5-20 μm) and permeability can be fine-tuned by adjusting the concentration and molecular weight of dextran. After encapsulation in these hydrogels, human mesenchymal stem cells and osteoblasts form a 3D cell network within 24 hours. The synthetic nature of this hydrogel enables histological analysis of cell-secreted collagen, a task previously challenging using collagen-derived hydrogels. Moreover, this hydrogel is integrated with a commercial chip, showcasing the potential for microfluidic perfusion cultures. Time-lapsed imaging of fluid flow and fast formation of 3D cell networks is demonstrated on chip. Altogether, this work introduces a versatile synthetic macroporous hydrogel, which can be integrated with microfluidic chip to enable 3D culture of human bone cell networks and analysis of cell-secreted ECM. This hydrogel may facilitate future mechanistic studies on bone development.

## 1. Introduction

Bone development — or osteogenesis — is a complex process that involves substantial changes in cell morphologies owing to a dynamic interplay between bone cells and their extracellular matrix (ECM).^[1]^ At an early stage of osteogenesis, osteoblasts lay down a thin layer of collagen-rich matrix (osteoid).^[2, 3]^ Thereafter, cells gradually embed themselves into this matrix to form a 3D cell network while inducing matrix mineralization. This process generates an intricate network of fluid-filled tunnels (i.e., lacunar-canalicular system), which have proven to be crucial for mediating load-induced bone formation.^[4-6]^

Although traditional bone tissue engineering centered on generating mineralized bone-like implants, there is a transition towards creating *in vitro* models of bone development for disease modeling and drug testing. Two major approaches have been sought: cell embedding in hydrogels and top seeding on porous scaffolds (Figure 1). For 3D embedding, a variety of natural (e.g., collagen,^[7]^ gelatin^[8]^) and synthetic hydrogels (e.g., clickable polyvinyl alcohol^[9]^ or poly(ethylene glycol) (PEG)^[10]^) have been reported. However, these hydrogels suffer from small pores (5-50 nm) and low permeability, which impede nutrient transport and cell-cell communication in 3D. In contrast, top seeding typically relies on macroporous scaffolds with large pores (100-600 μm)^[11-14]^ where the cell to surface interface is 2D. As a result, cells often fail to form 3D networks. Various techniques, such as emulsification^[15]^, porogen leaching^[16-18]^ and particle annealing^[19]^ have been employed to create macroporous hydrogels with relevant pore sizes. Nevertheless, these methods have limitations in generating interconnected pores in the presence of living cells.

**Figure 1.**
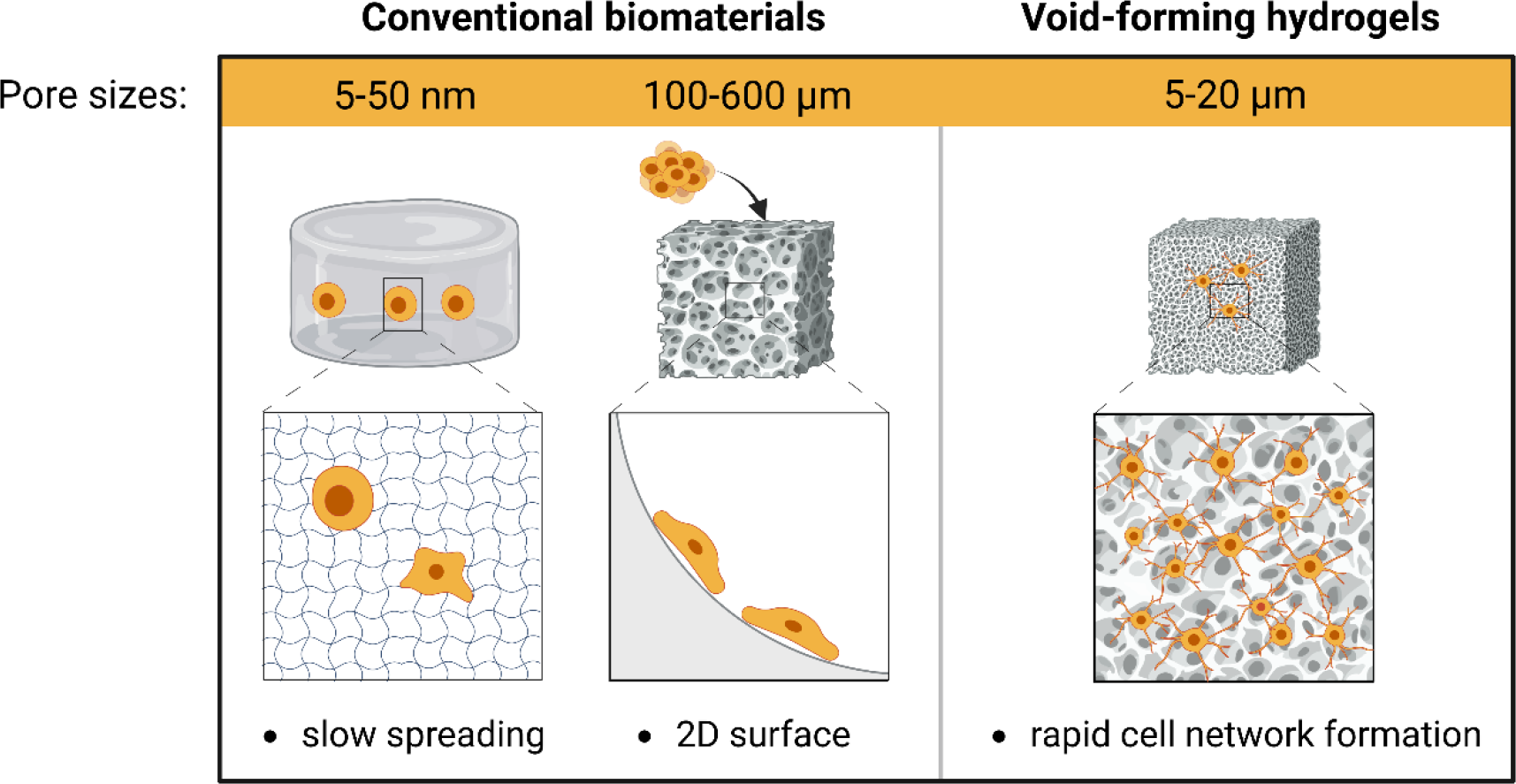
Schematic illustration of conventional biomaterials versus void-forming polyethylene glycol (PEG) hydrogels for bone tissue engineering. Traditional nanoporous hydrogels (pore sizes: 5-50 nm) impede cell spreading, while macroporous scaffolds with large pores (pore sizes: 100-600 μm) merely provide a 2D cell surface interface. Herein, void-forming PEG hydrogels (pore sizes: 5-20 μm) are developed for rapid 3D cell network formation. Illustration created with BioRender.com.

In the last decade, microfluidic organs-on-a-chip technology found increasing applications in tissue engineering.^[20-22]^ These tools provide exquisite control over environmental cues, such as mechanical and biochemical stimuli. Efforts to create bone-on-chip models have been actively sought for. For instance, Nasello et al.^[23]^ reported a static microfluidic culture of primary human osteoblasts in a collagen matrix. Notably, higher cell seeding density promoted osteogenic differentiation. Yet, using collagen limits the analysis of cell-secreted ECM, an important hallmark of bone development. Very recently, Bahmaee et al.^[24]^ combined porous polymerized high internal phase emulsion with a custom chip for perfusion culture of progenitor cells. To date, however, cells in current models fail to form a functional 3D cell network.

Herein, we report a synthetic biodegradable void-forming hydrogel for *in vitro* generation of functional 3D human bone cell networks and microfluidic integration. The synthetic nature of this hydrogel enables histological analysis of cell-secreted ECM during *in vitro* osteogenic differentiation. Optimization of pore size and permeability allows for successful integration within a microfluidic chip, time-lapsed flow imaging and proof-of-concept perfusion. This study introduces a synthetic hydrogel with fine-tuned microarchitecture for encapsulating living cells and generating functional cell networks to mimic early bone development *in vitro*.

## 2. Results and Discussion

### 2.1 Design of Void-Forming PEG Hydrogels

In the present work, a synthetic void-forming hydrogel was designed to generate 3D bone cell networks to mimic the process of osteoblast embedding during early osteogenesis. PEG hydrogels were formed by thiol-Michael crosslinking^[25]^ between 4-arm PEG vinylsulfone (4-PEG-VS) and di-thiol crosslinkers in the presence of dextran and hyaluronic acid (HA). This composition resulted in *in situ* pore formation by polymerization-induced phase separation (PIPS, Figure 2a). A fibronectin-derived arginylglycylaspartic acid peptide (N-C: CG**RGD**SP) was added to promote cell attachment, whereas a matrix metalloproteinase (MMP)-sensitive di-cysteine peptide^[9, 26]^ (N-C: KCGPQG↓IWGQCK or GCRDGPQG↓IWGQDRCG, ↓ indicates cleavage site) and PEG di-thiol^[27]^ (PEG-2-SH, *M*_*W*_=2.0 kDa or 3.4 kDa) were used as degradable and non-degradable crosslinkers, respectively. Using fluorescently labeled 4-PEG-VS, dynamic *in situ* pore formation was evidenced by time-lapsed confocal microscopy. After 1 h of crosslinking, spatial patterns of binodal nucleation were observed (Figure 2b, Supplementary Movie 1). Supplementary Figure 1 and Supplementary Movie 2 show that PIPS and interconnected porosity was only observed in the phase-separated hydrogels using dextran.

**Figure 2.**
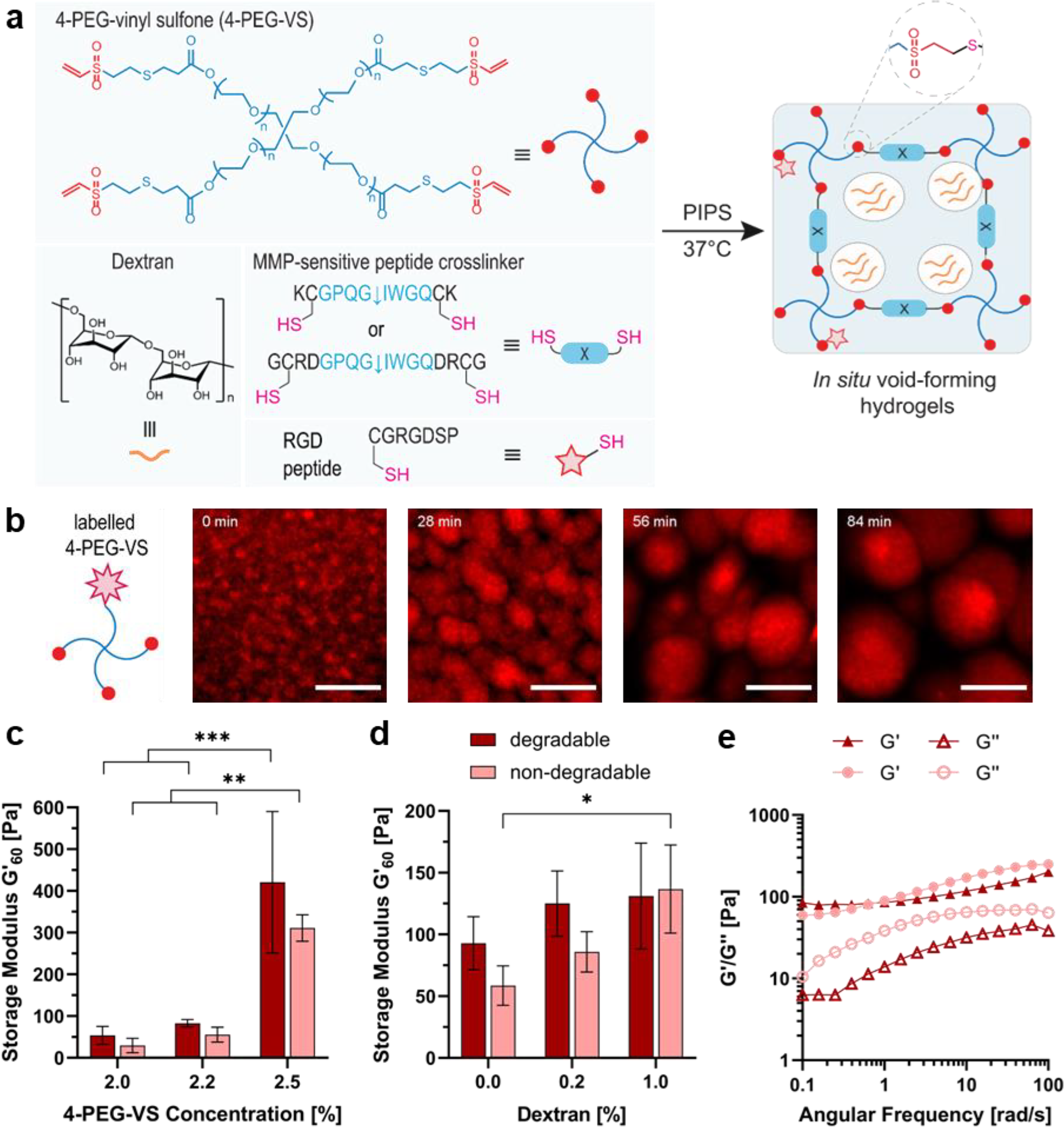
Synthesis and characterization of macroporous PEG hydrogels by polymerization-induced phase separation (PIPS). **a)** Illustration of *in situ* pore formation by PIPS at 37°C: upon addition of a matrix metalloproteinase (MMP)-degradable di-cysteine crosslinker, 4-arm PEG vinyl sulfone (4-PEG-VS) is crosslinked in the presence of dextran and hyaluronic acid (HA), leading to *in situ* pore formation. **b)** Time-lapsed confocal microscopy images showing PIPS between rhodamine-labeled 4-PEG-VS and dextran, scale bar: 10 μm. **c)** Storage modulus (*G’*) of macroporous hydrogels with varying 4-PEG-VS concentration and MMP-degradable peptide or PEG di-thiol as non-degradable crosslinker after 60 min of crosslinking at 37°C, *n*=3 (\*\**p* <0.01, \*\*\**p* <0.001, two-way ANOVA/Tukey). **d)** *G’* of degradable and non-degradable macroporous hydrogels with varying dextran concentration after 60 min of crosslinking at 37°C, n≥3 (\**p* <0.05, two-way ANOVA/Tukey). **e)** Viscoelasticity of macroporous PEG hydrogel matrices as determined by frequency sweep measurements on a rheometer at 37°C (oscillatory strain: 5%, angular frequency: 0.1–100 rad s^-1^).

### 2.2 Effect of Polymer Composition on Hydrogel Mechanics

Rheological analysis revealed that increasing the concentration of 4-PEG-VS from 2.0% to 2.5% yielded significantly stiffer hydrogels for both degradable and non-degradable groups with storage moduli (*G’*) ranging from 30 ± 17.2 Pa to 421 ± 169.5 Pa (Figure 2c). Depending on the hydrogel composition, gelation (crossover of *G’* and loss modulus (*G’’*)) started immediately or after up to 15 minutes upon *in situ* crosslinking at 37°C (Supplementary Figure 2a), whereby MMP-degradable hydrogels crosslinked faster than their non-degradable counterparts. Lutolf et al.^[28]^ demonstrated that the presence of positively charged amino acid residues near the cysteine of a crosslinker, such as the lysine residue in the MMP-sensitive crosslinker, modulates the thiol group’s pKa, thereby accelerating crosslinking kinetics in Michael-addition PEG hydrogels.

By increasing the dextran concentration from 0.0% to 1.0%, *G’* increased for both non-degradable and degradable hydrogels. This effect can be attributed to the strong phase-separating properties of dextran against PEG precursors, which may lead to a localized densification of the PEG phase and stiffer hydrogels. Varying the dextran molecular weight (*M*_*W*_, 40 kDa or 500 kDa), however, did not significantly affect the hydrogel mechanical properties (Supplementary Figure 2b). By changing the concentration of HA and thereby the viscosity of the hydrogel precursor, the stiffness of the hydrogel could be tuned (Supplementary Figure 2c). High viscosity has been suggested to prevent the phases from collapsing into microspheres before the structures are stabilized by crosslinking in PIPS.^[27]^ Our findings show that the inclusion of HA enhanced the crosslinking when its concentration was increased from 0.25% to 0.50%. However, higher concentration of HA (0.83%) reduced *G’*, indicating that crosslinking was less efficient for highly viscous formulations. Interestingly, a rheological test evidenced the viscoelasticity in the PEG hydrogel matrices for both degradable and non-degradable groups, presumably due to the HA content (Figure 2e). Qiu et al.^[29]^ reported that a viscous macromolecular component in a hydrogel can contribute significantly to the stress-relaxation properties and fast cellular morphogenesis. This property may facilitate fast 3D cellular network formation with PEG hydrogels.

### 2.3 Characterization of Pore Architecture

We investigated the impact of dextran inclusion on pore architecture in PEG hydrogels. The porosity after PIPS was analyzed by quantification of confocal microscopy images using an algorithm reported by Vandaele et al.^[30]^ As shown in Figure 3a-c and Supplementary Figure 3, the increase of dextran concentration from 0% to 0.5%, 1% and 2% led to an increase in pore radius (median pore radius: 4.0, 7.0, 10.0 and 10.0 μm, respectively) and porosity (25.1 ± 2.1, 37.6 ± 4.1, 50.4 ± 1.9 and 55.8 ± 5.9%, respectively) as well as higher pore connectivity. Furthermore, the increase of dextran *M*_*W*_ from 40 kDa to 500 kDa resulted in a shift towards larger pore radii (Figure 3d-e, Supplementary Figure 4). The resultant porosity in 500 kDa dextran group (19.7 ± 3.0%) was smaller than that of the 40 kDa group (24.7 ± 6.4%) but was higher than in compositions without the addition of dextran (12.0 ± 10.1%) (Figure 3f). Furthermore, we assessed the effect of physical constraints on pore formation (Supplementary Figure 5) inside a microfluidic channel. Similar as in non-constrained hydrogels, the pore radius for the high *M*_*W*_ dextran (500 kDa) group was larger (median: 4.2 μm) than that of the low *M*_*W*_ dextran (40 kDa) group (median: 2.6 μm). Moreover, pore connectivity on-chip was increased with higher dextran *M*_*W*_. These results demonstrate that the pore architecture of PEG hydrogels can be fine-tuned by dextran concentration and *M*_*W*_. The injectability of these synthetic hydrogels, along with their ability to retain void-forming characteristics when casted on a microfluidic chip, holds promise for microfluidic cell culture applications.

**Figure 3.**
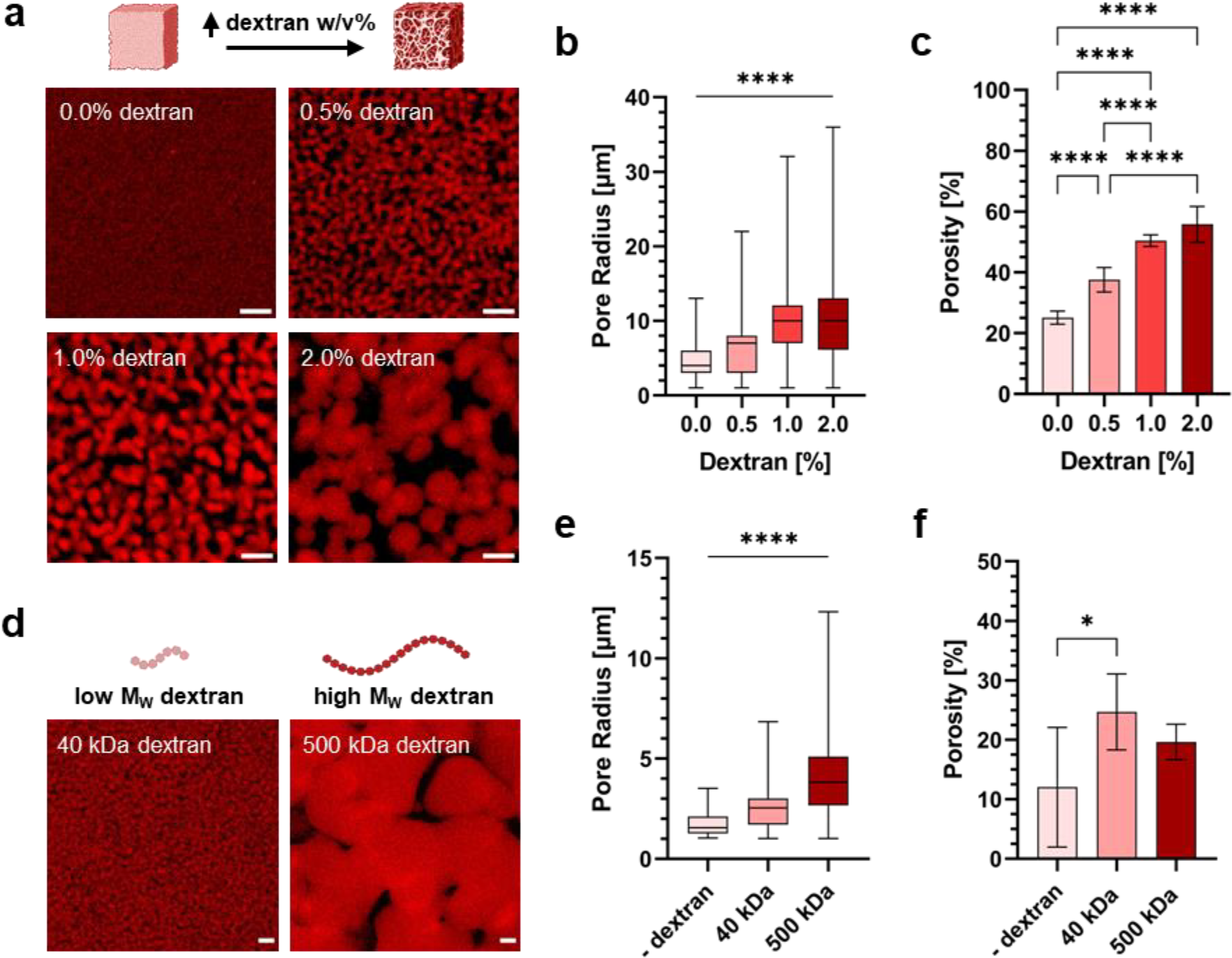
Characterization of the porous architecture of void-forming PEG hydrogels in function of dextran concentrations and molecular weights. **a)** Confocal microscopy images of rhodamine-labeled PEG hydrogels formed with varying dextran concentrations without physical constraints, scale bars: 10 μm. **b-c)** Quantification of pore radius and porosity of hydrogels formed with varying dextran concentrations, *n*=3 (\*\*\*\**p* <0.0001, one-way ANOVA/Tukey). **d)** Confocal microscopy images of rhodamine-labeled PEG hydrogels formed with 1.0% low *M*_*W*_ (40 kDa) and high *M*_*W*_ (500 kDa) dextran without physical constraints, scale bars: 10 μm. **e-f)** Quantification of pore radius and porosity of hydrogels formed with low *M*_*W*_ (40 kDa) and high *M*_*W*_ (500 kDa) dextran, *n*=3 (*p<0.05, \*\*\*\**p* <0.0001, one-way ANOVA/Tukey). Illustrations created with BioRender.com.

### 2.4 Rapid Formation of 3D Bone Cell Networks and Functional Maturation in a Static Culture

Previous studies have demonstrated the osteogenic potential of human mesenchymal stem cells (hMSCs) in void-forming hydrogels both *in vitro* and *in vivo*.^[16]^ In this study, we investigated whether our void-forming PEG hydrogels enable *in vitro* generation of 3D bone cell networks. hMSCs were embedded inside MMP-degradable and non-degradable PEG hydrogels and differentiated under osteogenic condition. MMPs are known to be crucial for hMSC differentiation as well as for osteoblast survival during osteogenesis.^[31]^ Therefore, we reason that only when cells are able to remodel their surrounding matrix through proteolysis by MMPs, they form an interconnected 3D cell network with long-term stability (Figure 4a). hMSC viability on day 2 was above 80% in both degradable and non-degradable hydrogels (Figure 4b). Notably, ultrafast cell spreading as well as 3D cell network formation within these macroporous hydrogels were respectively observed as early as 1.5 h and 24 h after being embedded in the hydrogel (Supplementary Figure 6). When embedded in MMP-sensitive hydrogels, hMSCs continuously remodel their environment and maintain a 3D cell network for at least 35 days. Quantification of mean cell area further evidenced the permissiveness of degradable hydrogels for hMSCs (Figure 4c). The average cell area in the degradable hydrogels was significantly larger compared to non-degradable hydrogels. An extensive 3D cell network formed in the degradable hydrogels on day 8. By contrast, the extent of cell network formation was significantly less in the non-degradable hydrogels (Figure 4d, Supplementary Figure 7). When grown in a more permissive environment, hMSCs formed multiple dendritic processes connecting them with neighboring cells.

**Figure 4.**
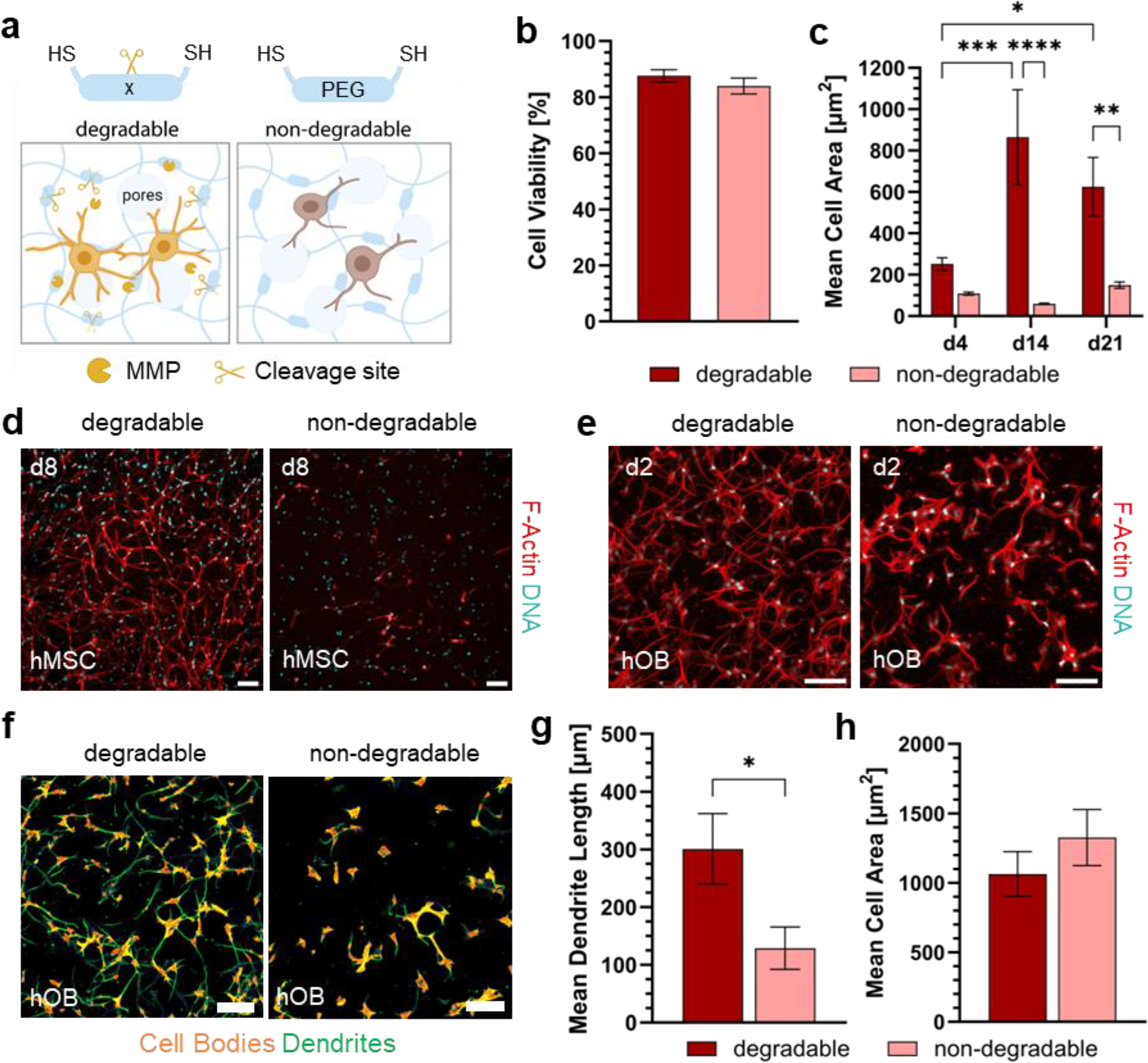
Static human bone cell culture within degradable and non-degradable void-forming PEG hydrogels. **a)** Illustration of the formation of 3D cell networks in degradable hydrogels (left) and the degeneration of cell networks in non-degradable hydrogels (right). Illustration created with BioRender.com. **b)** The impact of hydrogel composition on human mesenchymal stem cell (hMSC) viability on day 2, *n*=3 (Student’s t-test). **c)** The impact of hydrogel composition on mean hMSC area after 4, 14 and 21 days of 3D osteogenic culture, *n*=3 (\**p*<0.05, \*\**p*<0.01, \*\*\**p*<0.001, \*\*\*\**p*<0.0001, two-way ANOVA/Tukey). **d)** Representative maximum intensity projections (MIPs) of actin-nuclei-stained hMSC networks in degradable and non-degradable hydrogels after 8 days of culture, scale bars: 50 μm. **e)** Representative MIPs of actin-nuclei-stained human osteoblast (hOB) networks in degradable and non-degradable hydrogels after 2 days of culture, scale bars: 100 μm. **f)** Semiautomatic labelling of cell bodies and dendrites by NeuriteQuant plugin in ImageJ for quantitative comparisons of cell morphologies, scale bars: 100 μm. **g)** Quantification of mean dendrite length per cell using NeuriteQuant in degradable and non-degradable hydrogels, *n*=3 (\**p*<0.05, Student’s t-test). **h)** Quantification of mean cell area in degradable and non-degradable hydrogels, *n*=3 (Student’s t-test).

Based on the success of hMSC cultures, we further tested the suitability of macroporous PEG hydrogels for 3D osteogenic culture of human osteoblasts (hOBs). Cell viability after embedding was above 90% in both degradable and non-degradable groups and remained high after 2 days of culture (Supplementary Figure 8). Similar to hMSCs, a difference in cell morphology between degradable and non-degradable hydrogels was observed (Figure 4e). Quantification of cell denticity using ImageJ NeuriteQuant^[32]^ (Figure 4f) shows that hOBs within degradable PEG gels have longer dendritic processes (Figure 4g) despite having similar mean cell areas (Figure 4h). Moreover, we observed that the addition of cell-adhesive RGD-peptides is critical for hOB spreading and network formation (Supplementary Figure 9). Notably, we demonstrated that manipulating the porous architecture by increasing dextran concentration and, consequently, pore size influenced hOB morphology (Supplementary Figure 10a). Cells within PEG hydrogels with larger pore sizes (1.0% dextran) displayed a significantly larger cell area (Supplementary Figure 10b) and longer dendrites (Supplementary Figure 10c) on the second day of osteogenic culture compared to hydrogels with smaller pore sizes (0.2% dextran). In summary, our findings underscore the importance of both matrix degradability and porous architecture in the formation of long-term stable 3D cell networks from both hMSCs and hOBs.

### 2.5 Histological Analysis of Static Osteogenic Cultures

As the osteogenic differentiation proceeded, the cell-laden hydrogels became increasingly opaque. As such, histological sections of the same cultures on day 8 (Supplementary Figure 11) and day 30 (Figure 5) were prepared to further compare cell morphology, collagen secretion, matrix mineralization and osteocalcin expression. Similar to the non-sectioned samples, results show that degradable hydrogels facilitated pronounced cell network formation on day 8 (Supplementary Movie 3) that remained stable for at least 30 days in contrast to non-degradable hydrogels. Collagen type I is the major ECM protein secreted by osteoblasts during bone formation.^[33]^ Picrosirius red-polarization imaging revealed the presence of cell-secreted collagen fibers on day 8, especially in the MMP-degradable hydrogels due to its permissiveness for cell-matrix remodeling. Importantly, the synthetic nature of the macroporous PEG hydrogel allowed for assessment of collagen secretion considering that all detectable collagen fibers were produced by embedded cells. This is unachievable in conventional proteinaceous hydrogels such as collagen type I^[23]^ and gelatin derivatives.^[34]^

**Figure 5.**
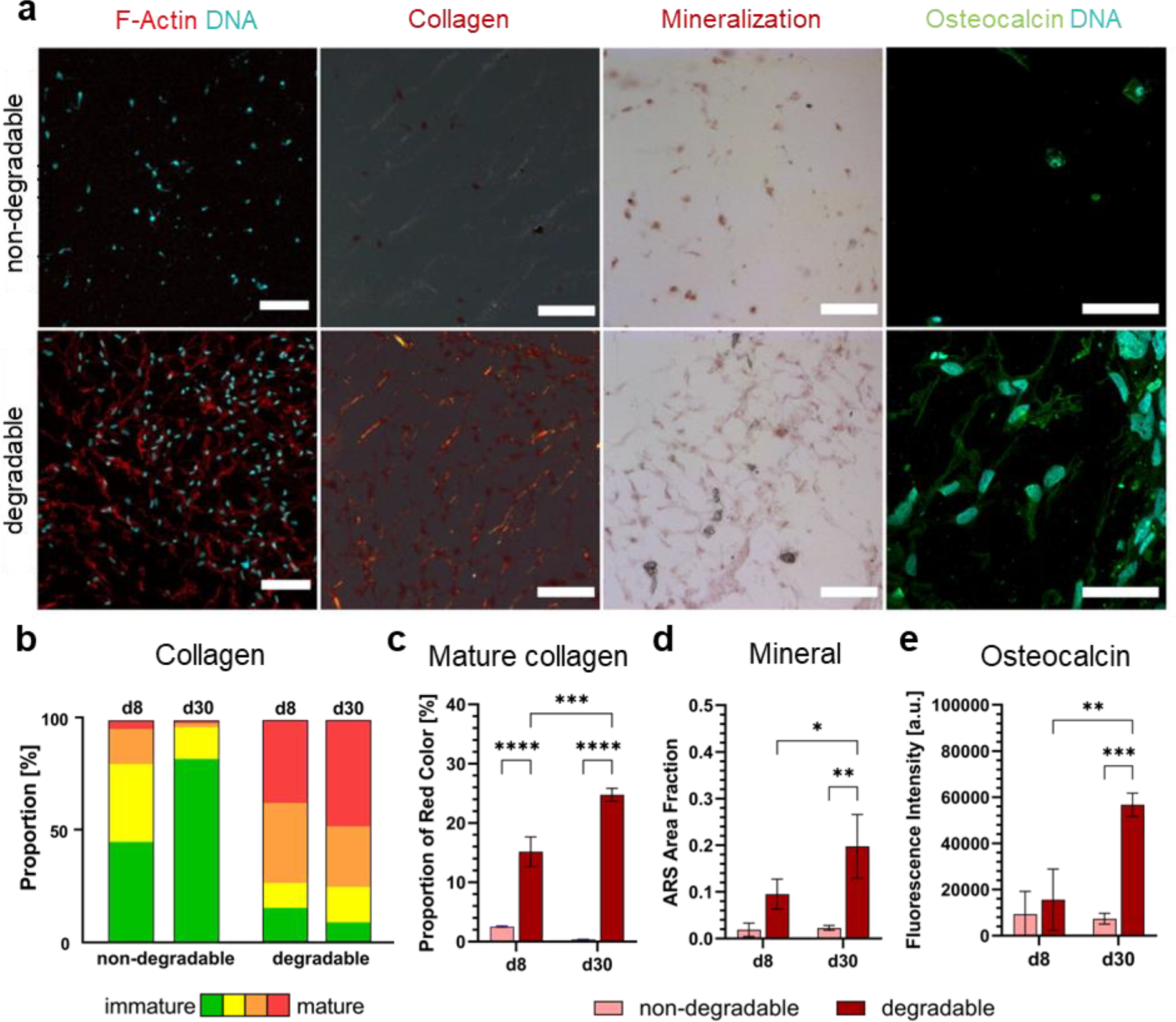
Histological analysis of static hMSC culture within MMP-degradable and non-degradable PEG hydrogels. **a)** Microscopy images of osteogenic markers on day 30, including cell morphology determined by confocal microscopy (MIP, scale bars: 100 μm), collagen fiber secretion determined by Picrosirius-polarization microscopy (scale bars: 100 μm), matrix mineralization determined by Alizarin red staining (scale bars: 100 μm) and osteocalcin expression by immunofluorescence staining (MIP, scale bars: 50 μm). **b)** Quantification of collagen content by fiber hue depending on the matrix degradability at day 8 and day 30, color indicates fiber thickness from green (thin, immature) to red (thick, mature), *n*=3. **c)** Quantification of red (mature) fiber content, *n*=3 (\*\*\**p*<0.001, \*\*\*\**p*<0.0001, two-way ANOVA/Tukey). **d)** Quantification of Alizarin red staining area fraction indicating the extent of matrix mineralization, *n*=3 (\**p*<0.05, \*\**p*<0.01, two-way ANOVA/Tukey). **e)** Quantification of osteocalcin expression by mean fluorescence intensity per cell, *n*=3 (\*\**p*<0.01, \*\*\**p*<0.001, two-way ANOVA/Tukey).

Collagen content and maturity were quantified based on the fiber hue method as described elsewhere.^[35]^ Figure 5b shows that more green and yellow color corresponding to low fiber thickness and immature collagen was present in the non-degradable hydrogels. In contrast, cells within the MMP-degradable hydrogels produced more mature collagen fibers as indicated by the larger proportion of red color (Figure 5c). Moreover, the red color content significantly increased from day 8 to day 30. Alizarin red staining further indicated enhanced matrix mineralization especially in close proximity to embedded cells within the degradable hydrogels compared to non-degradable ones after 30 days (Figure 5d). Compared to day 8, an increase in mineral deposition on day 30 implies the 3D osteogenic differentiation of hMSCs into a more mature bone cell phenotype. Osteocalcin, a marker for mature osteoblasts,^[36]^ was predominantly expressed in MMP-degradable hydrogels after cultivation for 30 days. In contrast, only limited expression of osteocalcin was observed in the non-degradable hydrogels (Figure 5e). Together, these results suggest that the MMP-degradable macroporous PEG hydrogels are permissive for the formation of 3D bone cell networks and allow for subsequent osteogenic differentiation in a static culture.

## 2.6 On-Chip Integration of Macroporous PEG Hydrogels

For potential microfluidic applications, we injected the void-forming hydrogels into a microfluidic chip and introduced perfusion through the porous matrix. Flow visualization showed that interstitial liquids could pass through the macroporous PEG hydrogels when applying a pressure gradient on a microfluidic setup (Figure 6a). The pressure difference across an acellular PEG hydrogel was created by applying dissimilar flow rates (*Q*_*1*_>*Q*_*2*_) to the two medium channels. The perfusion of different fluorescein isothiocyanate (FITC)-dextran tracer molecules (70 kDa and 500 kDa) was visualized by fluorescence microscopy. Time-lapsed microscopy images (Supplementary Movie 4) indicate that fluorescence intensity increased gradually for both tracers over time. The intensity of tracer molecules increased until reaching a plateau (Supplementary Figure 12). Yet, no statistical difference in the intensity change was observed between the two tracer molecules. We thus reason that pores within PEG hydrogels were large enough for both sizes of tracer molecules to pass through. These results correlate with the pore size quantification (Figure 3) that pore sizes in PEG hydrogels are in the order of μm, which are much larger than the sizes of tracers.

**Figure 6.**
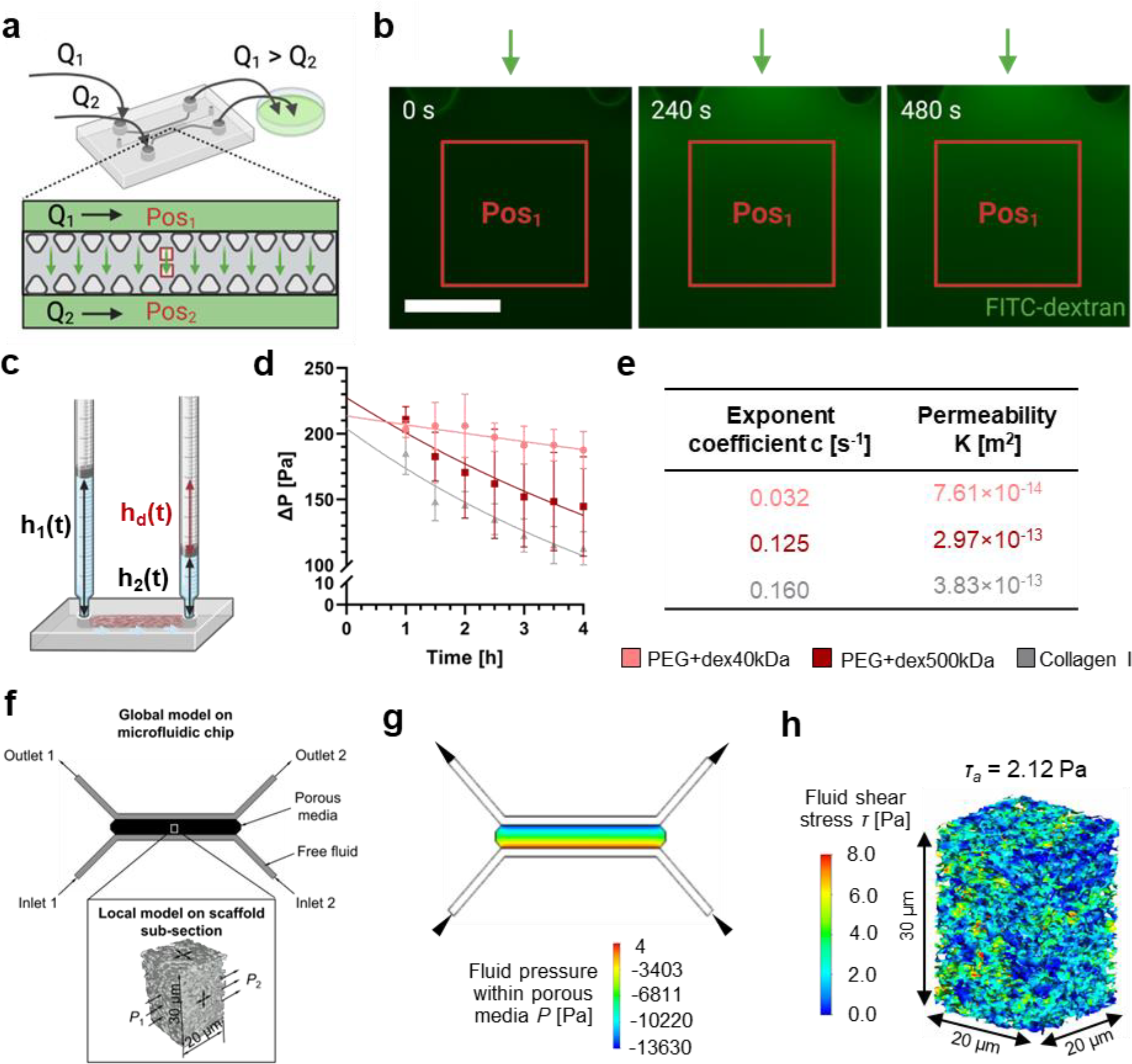
Integration of PEG hydrogels with a microfluidic chip for perfusion of the macroporous matrix. **a)** Microfluidic setup to visualize fluid flow through PEG hydrogels, *Q*_*1*_ and *Q*_*2*_ denote volumetric flow rate, Pos_1_ and Pos_2_ represent positions of image acquisition. **b)** Time-lapsed fluorescence microscopy images of fluorescein isothiocyanate (FITC)-dextran (500 kDa) tracer perfusing through PEG hydrogels on chip in response to a pressure gradient, scale bar: 200 μm. **c)** Microfluidic setup for permeability test depending on a gravity-driven fluid flow through the PEG hydrogels, *h*_*1*_ and *h*_*2*_ denote measured heights, *h*_*d*_ is the height difference to calculate the changes in pressure drop over time. **d)** Calculated pressure change over time (symbols) and fitted exponential functions (lines) across 40 kDa and 500 kDa dextran PEG hydrogels and reference hydrogels made of 2 mg ml^-1^ collagen type I, *n*=3. **e)** Resulting Darcy’s permeability of the 3 hydrogels obtained from pressure change over time. **f)** Global multiphasic computational fluid dynamics (CFD) model of microfluidic device with the whole scaffold (porous media domain) in it and local CFD model of the subsection whose struts geometry is constructed from confocal images of fluorescently labeled PEG hydrogel formed inside the microfluidic chip. **g)** Pressure distribution within the porous media (homogenized scaffold domain) under the applied flow rate of 10 μl min^-1^ per inlet. **h)** Fluid shear stress (FSS) distribution and average FSS (*??*_*a*_) within a representative subsection (x-y-z: 20 x 20 x 30 μm) under an applied flow rate of 10 μl min^-1^ per inlet. Illustration created with BioRender.com.

To quantify hydrogel permeability, we employed a microfluidic setup (Figure 6c) to create a pressure difference across the hydrogel caused by different volumes of medium added to each syringe barrel. The change in pressure difference over the hydrogel was monitored and an exponential decay function was fitted (Figure 6d). The values for exponent coefficient *c* (in the equation in Supplementary Method – *Quantification of Permeability*) and calculated Darcy’s permeability *K* are summarized in Figure 6e. The addition of 500 kDa dextran increased the permeability of the macroporous hydrogel compared to 40 kDa dextran as expected given the larger pore sizes as shown in Figure 3. Importantly, the permeability is comparable to that of a reference hydrogel made of collagen type I. We envisage that these void-forming PEG hydrogels hold great potential for future on-chip perfusion cultures using a chemically defined macroporous hydrogel.

Next, we applied a multiscale and multiphase computational fluid dynamics (CFD) model^[37]^ to estimate the fluid shear stress (FSS) within the macroporous hydrogel (Figure 6f). Using the permeability values of macroporous PEG hydrogels with 40 kDa dextran, a global model on the microfluidic chip was generated to estimate the pressure gradient across the hydrogel region by applying two different flow rates (*Q*_*low*_=10 μl min^-1^ and *Q*_*high*_=100 μl min^-1^ per inlet, Figure 6g). According to CFD calculations, the pressure drop over a hydrogel subsection in the center of the channel at *Q*_*low*_ was 196 Pa. This value was used for defining the loading conditions of the local model as illustrated in Figure 6f. After solving the local CFD model of 4 discretized subsections, the FSS distribution of a representative subsection is shown in Figure 6h. The overall average FSS (*τ*_*a*_) at *Q*_*low*_ was 1.74 Pa. Considering the FSS is proportional to the applied flow rate, the average FSS at *Q*_*high*_ was 17.43 Pa. These results imply that the majority of the FSS values inside the porous geometry at *Q*_*low*_ is within the physiological range for bone tissues,^[38]^ whereas it is far exceeded at *Q*_*high*_.

The estimation of FSS within macroporous PEG hydrogels during perfusion serves as a valuable approach to approximate the mechanical cues experienced by embedded cells under dynamic culture conditions. In this study, we used this estimation as an initial step for a preliminary dynamic bone cell culture on a chip. However, it is essential to note that this model does not account for the impact of embedded cell networks on hydrogel permeability or the complexities of dynamic cell-matrix remodeling, which can alter fluid dynamics. Moreover, the FSS was calculated based on permeability values from PEG hydrogels with 40 kDa dextran and would be different when using more permeable 500 kDa dextran. To gain a more comprehensive understanding of how FSS affects bone cells in microfluidic culture, more advanced CFD models are therefore warranted for future investigations.

### 2.7 On-Chip Culture of 3D Bone Cell Networks

We investigated the feasibility of generating 3D bone cell networks on a chip. Figure 7a shows hMSCs embedded in both degradable and non-degradable matrices during the second day of osteogenic culture. Remarkably, similar to off-chip cultures, hMSCs exhibited remarkable capacity for rapid spreading, forming intricate 3D cell networks with elongated dendritic protrusions after two days, within the degradable matrix.

**Figure 7.**
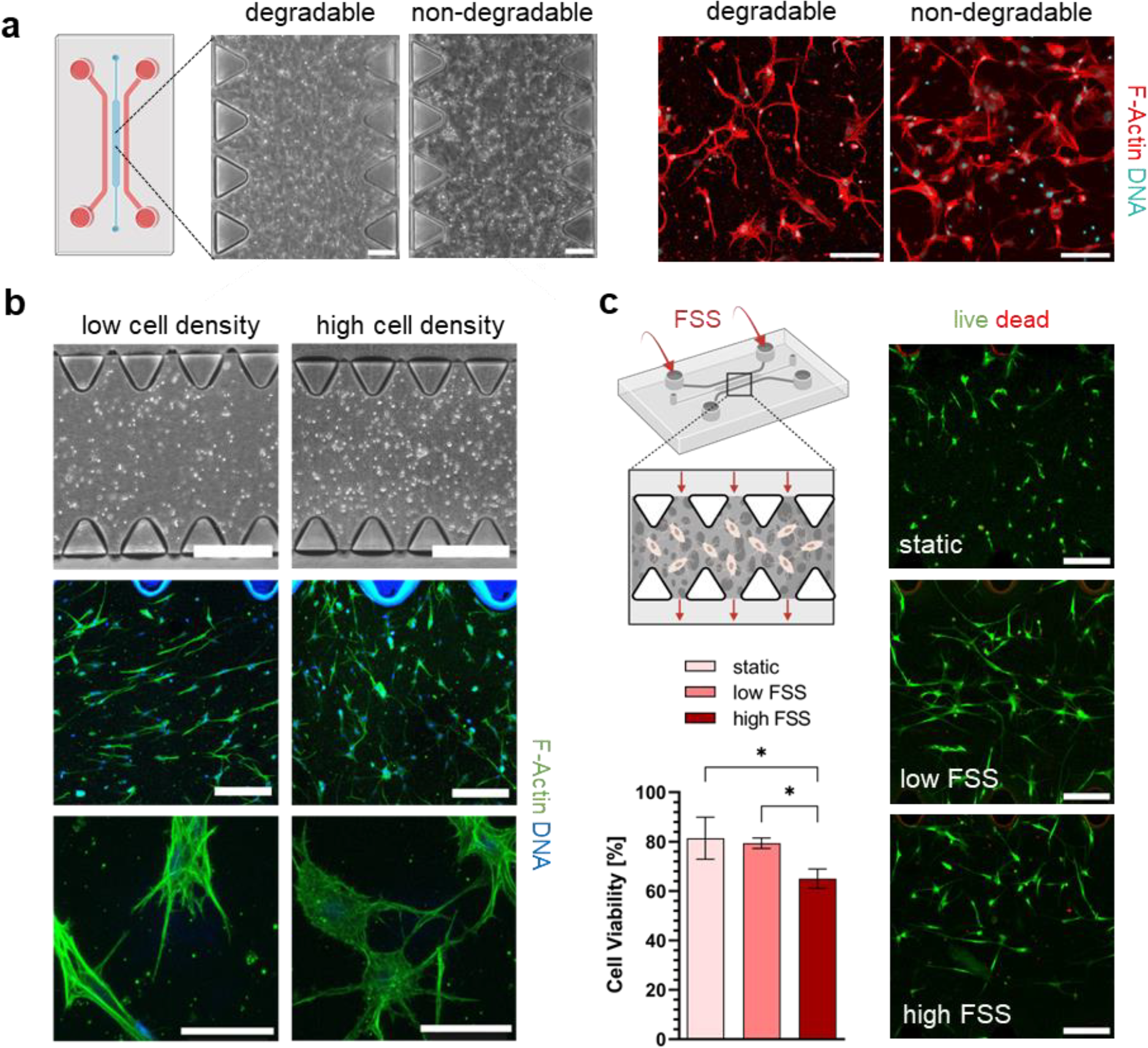
3D microfluidic culture of hMSCs within PEG hydrogels. **a)** Phase contrast (scale bars: 200 μm) and confocal microscopy images (scale bars: 100 μm) showing cell network formation of hMSCs embedded in degradable and non-degradable PEG hydrogels on chip after 2 days of osteogenic culture. **b)** Phase contrast and confocal microscopy images showing the effect of high (1×10^6^ ml^-1^) and low (5×10^5^ ml^-1^) cell seeding density on hMSC morphology inside MMP-degradable PEG hydrogels on day 7 of osteogenic culture on chip, scale bars: top 500 μm, middle - 200 μm, bottom - 50 μm. c) Cell viability on day 13 of hMSCs embedded in MMP-degradable PEG hydrogels on chip in response to low FSS (20 μl min^-1^) or high FSS (200 μl min^-1^) or static culture, *n*=3 (\**p*<0.05, one-way ANOVA/Tukey), scale bars: 200 μm.

Cell seeding density has been shown as a key factor in dictating 3D differentiation of primary hOB within a collagen type I hydrogel.^[23]^ We further studied how different cell seeding density (low: 5×10^5^ ml^-1^ and high: 1×10^6^ ml^-1^) influences the formation of 3D bone cell networks in degradable macroporous PEG hydrogels on a chip (Figure 7b). Following seven days of cultivation in osteogenic medium, hMSCs cultivated at high seeding density exhibited a dendritic morphology, actively forming an intricate 3D cell network. Conversely, cells at low seeding density appeared elongated, yet failed to establish intercellular connections.

Given their macroporosity, permeability and permissiveness for differentiation, we lastly examined the potential of void-forming PEG hydrogels for dynamic cell culture on a chip. Specifically, we studied the effect of controlled delivery of FSS on the viability of embedded hMSCs. Cells were subjected to mechanical stimulation for 2×10 minutes per day, following the same regimen as the CFD model, using either a low flow rate (*Q*_*low*_=10 μl min^-1^ per inlet) or high flow rate (*Q*_*high*_=100 μl min^-1^ per inlet), while cells cultivated under static conditions were selected as the control (Figure 7c). The live/dead assay results on day 13 revealed comparable cell viability between static and low FSS culture, while high FSS led to a significant reduction in cell viability. The cell death under high FSS could be attributed to high shear stresses well beyond the physiological range. Indeed, previous studies have shown that laminar flow at physiological levels exerts an anti-apoptotic effect on MSCs^[39]^ and osteocytes.^[40]^ The approximate average FSS *τ*_*a*_ at *Q*_*high*_ (17.43 Pa) calculated by the CFD simulation far exceeded the physiological range within the LCN porosities (0.8–2.0 Pa)^[38]^, resulting in more apoptotic cells compared to the static and low FSS groups, where the average FSS *τ*_*a*_ was estimated to be 1.74 Pa. This preliminary investigation highlights the potential of void-forming PEG hydrogels for dynamic cell culture on a chip and sheds light on the influence of FSS on hMSC viability. However, to establish definitive conclusions regarding the effect of perfusion on osteogenic differentiation, in-depth dynamic perfusion cultures are warranted.

In conclusion, we have successfully developed a synthetic macroporous PEG hydrogel that effectively supports 3D culture of human bone cell networks from primary cells. By adjusting the mechanical and microarchitectural properties, as well as the degradability of the hydrogel, rapid cell spreading and osteogenic differentiation are promoted. Within the MMP-degradable matrices, single bone cell precursors respond to the porous architecture, forming an interconnected cell network in 3D without adding morphogens^[41]^, and differentiating into an osteoid-like tissue. An essential advantage of this technique lies in the synthetic nature of these hydrogels, which are chemically defined and have minimal batch-to-batch variations. Furthermore, the use of this synthetic non-fibrillar hydrogel facilitates histological analysis of cell-secreted collagen fibers as a potential biomarker for studying (patho-)physiological processes linked to bone formation. Moreover, the unique process of PIPS allows for the formation of interconnected pores in the presence of living cells, a feature unachievable with other types of macroporous hydrogels formed through previously described methods.^[15-17, 19]^ The resulting permeability allows for fluid flow through the matrix, making it compatible with microfluidic platforms and the potential to deliver mechanical stimuli to enhance osteogenesis. This *in vitro* platform may open up new avenues for future mechanistic studies on bone development, providing valuable insights into both healthy and pathological conditions.

## 4. Experimental Section

### Hydrogel Preparation

To prepare (labeled) PEG hydrogels, (rhodamine-labeled) 4-PEG-VS (20% w/v in HEPES), 40 kDa or 500 kDa dextran (Sigma-Aldrich, 31389-25G or 31392-10G, in phosphate buffered saline (PBS) pH 7.4), RGD (China Peptides, N-C: CG**RGD**SP, in PBS pH 7.4 or pH 6), HA (Sigma-Aldrich, 9067-32-7, 0.5%–1% w/v in Hanks’ Balanced Salt Solution (Gibco, 14025-050) or Dulbecco’s modified Eagle’s medium (DMEM, Gibco, high glucose)) and crosslinker (PEG-2-SH (2 kDa or 3.4 kDa, Laysan Bio, in PBS pH 7.4 or pH 6) or MMP-degradable (China Peptides, N-C: KCGPQGIWGQCK or GCRDGPQGIWGQDRCG, in PBS pH 7.4 or pH 6 or TEOA)) were thoroughly mixed in this order to ensure PIPS, efficient crosslinking and even distribution of RGD. RGD and crosslinker stock solutions were prepared directly before mixing the precursor solution and kept on ice to prevent rapid oxidation of thiol groups. Final concentrations of each component are noted in the figure captions. Typical ranges were 2.0–2.5% w/v 4-PEG-VS, 0.0–2.0% w/v dextran, 0.25–0.83% w/v HA and a thiol/ene ratio of 0.8 or 1.25 between 4-PEG-VS and crosslinker and 0.07 between 4-PEG-VS and RGD. The exact composition for each experiment is shown in Supplementary Table 1. Once the hydrogel precursor solution was prepared, it was rapidly casted into either a custom-made round poly(dimethylsiloxane) (PDMS) mold on a confocal dish (VWR, 734-2905) or into the central hydrogel channel of a 3D cell culture chip (AIM Biotech, DAX-1) inside the corresponding chip holder (AIM Biotech, HOL-1 or HOL-2) according to the manufacturer’s protocol. Hydrogels were crosslinked at 37°C and 5% CO_2_ for at least 60 min. After crosslinking, hydrogels were washed 1–2 times with PBS.

### Characterization of Porosity

For a quantitative analysis of pore size and pore connectivity, hydrogels with rhodamine-labeled 4-PEG-VS were prepared, washed with PBS, and incubated at 37°C to allow for swelling for at least 24h. Hydrogels were then imaged by confocal microscopy. Z-stacks of 20–30 μm were obtained and deconvoluted using Huygens Professional Software. These images were then processed with the MATLAB algorithm developed by Vandaele et al.^[30]^ that uses image segmentation and approximation of pores by spheres to quantify pore size, connectivity and other determinants of the porous architecture. An adaptive threshold sensitivity of 0.6 was chosen. Pore size (external pore radius in 3D), pore connectivity (number of neighboring pores) and porosity were obtained. Pores with external radii above 1 μm were considered for quantification.

### Quantification of Cell Morphology

To quantitatively compare mean cell area between groups, confocal microscopy z-stacks of 100–200 μm were acquired with a 25× objective. Using the MIP of these images, mean cell area was calculated by determining the area of actin in Fiji/ImageJ and dividing it by the number of nuclei in the same image. To analyze the dendrite length of cells embedded within macroporous hydrogels, MIPs (100 μm) of confocal microscopy images of actin-stained samples were used. Images were processed using the ImageJ plugin NeuriteQuant^[32]^ to segment cells into cell bodies and dendrites and subsequently quantify the mean length of all dendrites of a single cell.

### Collagen Imaging and Quantification

Cell-secreted collagen was investigated using Picrosirius red staining (Sigma-Aldrich, 365548). Briefly, the cryosections were stained in picrosirius red (0.1% in saturated aqueous picric acid) for 1 h and washed in two changes of acidified water. Polarized light microscopy images (polarization filter angles at 0° and 90°) were taken in transmission mode with a Zeiss AxioImager.Z2 at 20× magnification. Collagen content was then quantified according to the method described by Rich and Whittaker.^[35]^ First, images were converted to 8-bit RGB. The color threshold in the Hue spectrum was next split into the different colors as: red 2–9 and 230–256, orange 10–38, yellow 39–51 and green 52–128. The pixel area of each color was measured. The different hue ranges measured were expressed as a percentage of all pixels in the image. Fiber thickness and maturity of collagen fiber increases from green, yellow, orange to red.

### Visualization of Fluid Flow on Chip

To visualize fluid flow through the porous matrix, the hydrogel was casted into a microfluidic chip (AIM Biotech, DAX-1) with attached luer connectors (AIM Biotech, LUC-1). After hydration for 24 h, flow imaging was performed on a wide field microscope (Olympus, IX83). Two different tracer solutions were prepared by diluting a 0.1% w/v stock solution of 70 kDa or 500 kDa FITC-dextran (both Sigma-Aldrich, FD70S-100MG and FD500S-100MG) 1:1000 in DMEM. The chips were then mounted onto the microscope stage, the tracer solutions were filled into 20 ml syringes which were connected to the luer connectors on the chips via needles (0.80×22 mm, blunt, Braun), tubing (Semadeni Plastics, 4348) and male luer connectors (Cole Parmer, 45518-07) that were primed with liquid to prevent air from being trapped along the flow path. Using 2 syringe pumps (WPI, AL-1000), an interstitial flow was created by applying different flow rates (*Q*_*1*_=334 μl min^-1^ and *Q*_*2*_=10 μl min^-1^) to the medium channels. The perfusion of the PEG hydrogel with tracer molecules was imaged in two different positions using a filter for FITC and a 20× air objective every 20 s for 8 min. The pumps were switched on between time-points 2 and 3. For quantification, the mean intensity in a rectangular area at positions Pos_1_ and Pos_2_ was measured in Fiji/ImageJ for each time-point and both tracers. This intensity was normalized by the mean intensity at time-point 0 for each tracer and position.

### Statistics

Statistical analysis was performed in GraphPad Prism 8.2.0. Depending on the number of variables, ordinary one-way or two-way analysis of variance (ANOVA) were used followed by Tukey’s test for multiple comparisons if more than two groups were compared. For the comparison of only two groups, Student’s t-test (two-tailed) was used. Further, p values are indicated by * (<0.05), ** (<0.01), *** (<0.001) and ****(<0.0001) in figures. Data is shown as mean ± SD and the number of replicates is indicated in the figure caption. Whiskers of boxplots indicate minimum and maximum values.

## Supplementary Information

Supplementary Information is available from the online version of the paper.

## Author Contribution Statement

X.H.Q., D.Z., M.Z.M. conceived the study and designed the experiments. D.Z. carried out most of the experiments except static long-term hMSC culture and histological analysis. X.H.Q. and M.Z.M. co-established the void-forming hydrogels and static culture. S.S.L. contributed microfluidic imaging technique and co-supervision. M.H. contributed to hOB culture and rheological analysis. L.B. performed histological analysis on static culture samples. F.Z. carried out the CFD simulations. R.M. contributed to scientific discussion and co-supervision. All authors analyzed and interpreted the results. D.Z. wrote the manuscript with input from all authors.

## Conflict of Interest

The authors declare no conflict of interest.

## Acknowledgements

This project was supported by the Swiss National Science Foundation (SNSF) (project no. 190345, 206501). The authors thank Sophie Zengerle, Isabel Hui, Wanwan Qiu and Christian Gehre for experimental support as well as Dr. Tobias Schwarz in the Scientific Center for Optical and Electron Microscopy (ScopeM) of ETH Zurich for technical support and the University Children’s Hospital Zurich for supplying primary human osteoblasts.

## Notes

### Competing Interest Statement

The authors have declared no competing interest.

