## Supplementary Information for "Synthetic Biodegradable Void-forming Hydrogels for *In Vitro* 3D Culture of Functional Human Bone Cell Networks"

Dr. F. Zhao
Department of Biomedical Engineering and Zienkiewicz Centre for Computational Engineering, Faculty of Science & Engineering, Swansea University, Bay Campus, Swansea, SA1 8EN, United Kingdom

Dr. S. S. Lee
Scientific Center of Optical and Electron Microscopy, Institute of Biochemistry, ETH Zürich, Otto-Stern-Weg 3, 8093 Zürich, Switzerland

### Supplementary Methods

*Synthesis of 4-arm poly(ethylene glycol) vinylsulfone (4-PEG-VS)*: Rhodamine-labeled 4-PEG-VS and 4-PEG-VS were synthesized according to the protocol by Broguiere et al.[1] In a 2-neck flask, 4 ml of triethanolamine (TEOA, Sigma-Aldrich, 90279- 100ML) buffer (200 mM, pH 8.0) was purged with argon for 15 min to remove oxygen. Then, 1 g (50 µmol) of 20 kDa 4-PEG-thiol (Laysan Bio, SH-20K-1g) was added and stirred under argon protection. After dissolution, the solution was carefully transferred into a syringe and immediately added dropwise over vigorous stirring to 1.05 ml (10 mmol) of divinyl sulfone (97%, Alfa Aesar, L12827-09) in 4 ml of the same buffer. The reaction was left to proceed for 2 h. The product was dialyzed ($M_W$ cut-off: 3.5 kDa) against ultrapure water with 6 changes of water over at least 4 h each, sterile filtered, and lyophilized after removal of excessive water using a rotary evaporator (Heidolph, 55 mbar, 200 rpm, 40°C). The product was aliquoted and stored at -20°C until use.

*Synthesis of Rhodamine-labeled 4-PEG-VS:* In a flask, 2 ml of ultrapure water was protected with argon for 15 min. Then, 200 mg (10 µmol) of 20 kDa 4-PEG-thiol was dissolved in it. To this flask, a solution containing 0.24 mg (0.5 µmol) of tetramethyl rhodamine-5-maleimide (Sigma-Aldrich, 94506) in 2 ml phosphate buffered saline (PBS, Gibco, 10010-015, pH 7.4) was added dropwise over stirring. The conjugation happened within seconds but was left to proceed for 10 minutes. The resulting mixture was added dropwise into 462.5 µl (4.4 mmol) of divinyl sulfone in 4 ml TEOA buffer (200 mM, pH 8.0), left to react for 60 min under stirring, dialyzed, sterile filtered, aliquoted, and lyophilized. All the handling was performed in the dark. This protocol substitutes 1/80th of the 4-arm-PEG ends in the upper limit case of 100% conjugation efficiency.

*Time-Lapsed Pore Imaging:* For time-lapsed imaging of polymerization-induced phase separation (PIPS), rhodamine-labeled 4-PEG-VS was used when preparing the hydrogel precursor solution to image the 4-PEG-VS phase in the hydrogel. The hydrogel was crosslinked directly in a custom-made poly(dimethylsiloxane) (PDMS) mold inside a confocal microscope (Leica SP8) at 37°C. Using a 63× oil immersion objective with 1.4× zoom, z-stacks of 35 µm were obtained every 2 min for 90 min.

*Rheology:* For rheology, acellular hydrogel precursor solutions with varying compositions were prepared as described above and analyzed on an Anton Paar rheometer MCR302 (82868246) using a PP20 plate and a glass bottom. Hydrogels were crosslinked at 37°C for 60 min while a time-sweep measurement was performed at 1 Hz, 5% strain and with a gap of 100 µm to determine the hydrogel storage modulus (*G′*). For each sample, 40–50 µl hydrogel solution was loaded into the center of the glass plate and after setting the PP20 plate to the desired gap position, mineral oil (Sigma-Aldrich, 330779) was placed around the hydrogel to prevent dehydration. Frequency sweep was performed to test time-dependent viscoelastic properties of the hydrogels (oscillatory strain: 5%, angular frequency: 0.1-100 rad $s^{-1}$, 37°C).

*Human Mesenchymal Stem Cell (hMSC) Culture:* For 2D cell expansion, hMSCs (Lonza, PT-2501) were cultured in expansion medium containing Dulbecco's modified Eagle's medium (DMEM, Gibco, high glucose) with 10% v/v fetal bovine serum (FBS, Gibco, Lot#42F7190K or 2440094), 1% v/v Antibiotic-Antimycotic (Anti-Anti, Gibco, 15240-062), 1% v/v non-essential amino acids (Gibco, 11140-035) and 1 ng $ml^{-1}$ basic fibroblast growth factor (Invitrogen, 13256-029) in T150 cell culture flasks (TPP, 90151) at 37°C with 5% $CO_2$ until reaching 80% confluency. Medium was exchanged 3 times per week. Cells were washed twice with PBS (37°C) before adding 0.25% trypsin-EDTA (Gibco, 25200-056) to detach them from the flask. Trypsin activity was blocked by adding control medium (DMEM + 10% v/v FBS + 1% v/v Anti-Anti). Cells were counted using a hemocytometer and resuspended in control medium at the desired concentration. hMSCs (p5–p8) were embedded inside macroporous hydrogels by resuspending them in the hyaluronic acid (HA) stock solution during hydrogel preparation to obtain final cell concentrations of $5\times10^5$–$5\times10^6$ ml$^{-1}$. They were subsequently cultured in osteogenic differentiation medium (control medium with 10 mM β- glycerophosphate (Acros, 410991000), 50 µg ml$^{-1}$ L-ascorbic acid (Sigma-Aldrich, A92902-100G) and 100 nM dexamethasone (Sigma-Aldrich, D2915)). Medium was replaced 3 times a week in custom PDMS molds and 5 times a week for static culture on chip. For dynamic cell culture, fluid shear stress (FSS) was applied on chip (AIM Biotech, DAX-1) by connecting two syringe pumps to both inlets of one medium channel and applying a total flow rate of 20 µl min$^{-1}$ (low FSS) or 200 µl min$^{-1}$ (high FSS) using DMEM. Loading was performed 2×10 min daily starting on day 3 until day 7 of culture and then again from day 10 until day 13 with 60 min in between each treatment. Three replicates were used per condition (static, low FSS, high FSS).

*Human Osteoblast (hOB) Culture:* Primary hOBs were obtained from a commercial supplier (PromoCell, C-12720) or from the University Children's Hospital Zurich under ethical approval (KEK-ZH-Nr. 2019-00811) from healthy donors. For 2D cell expansion, hOBs (p5–p6) were cultured similarly as hMSCs until reaching 80% confluency. After embedding in PEG hydrogels, hOBs were cultured in the same osteogenic differentiation medium as hMSCs for 2 days before fixation.

*Live/Dead Assay:* To quantify cell viability, staining with Calcein Green AM (CaAM, Sigma-Aldrich, 56436-50UG) and Ethidium-homodimer-1 (EthD-1, Sigma-Aldrich, 460439) was performed. Staining solution (1:1000 EthD-1 and 1:500 CaAM in PBS) was applied after washing samples twice with PBS and then incubated for 15–20 min at 37°C protected from light before washing again with PBS. Samples were imaged using confocal microscopy with a 10× air objective. For analysis, maximum intensity projections (MIP) of z-stacks of 70–100 µm each were created in Fiji/ImageJ. Cells in green and red channel were either counted manually if discrimination between single cells was not possible or a custom-written macro was used. Viability was then calculated as the percentage of live cells among all present cells in the MIP.

*Fixation and F-Actin-Nuclei Staining:* At the end of culture, cells were fixed by first washing them with PBS and then applying a solution of 4% paraformaldehyde (Sigma-Aldrich, 15-812-7) for 15–20 min at room temperature. Samples were washed twice with PBS. F-Actin-nuclei staining was performed to further investigate cellular morphology and cell network formation. Hydrogels were incubated in 1% v/v bovine serum albumin (BSA, Sigma-Aldrich, 9048-46-8) in PBS for 1.5 h at room temperature. Subsequently, cells were permeabilized in a solution of 0.2% w/v Triton X-100 (Sigma-Aldrich, 9002-93-1) in 0.1% BSA in PBS for 10 min. Hydrogels were washed 3 times with PBS. The staining solution containing dilutions of 1:1000 Hoechst 33342 (1:1000, Sigma-Aldrich, B226) and 1:200 Phalloidin CruzFluor 647 Conjugate (Santa Cruz Biotechnology, sc-363797) or Phalloidin-TRITC (Sigma-Aldrich, P1951) in 0.1% BSA was prepared. On-chip samples were stained for 12–24 h at 4°C, samples in confocal dishes for 1.5–2.0 h at room temperature protected from light. Before image acquisition, on-chip samples were washed 5 times with 5 min between each wash and hydrogels on confocal dishes were washed 3 times. Imaging was performed using confocal microscopy.

*Cryosectioning:* Prior to cryosectioning, fixed samples were cryoprotected overnight in 30% w/v sucrose (Sigma-Aldrich, S7903) in PBS at 4°C. The next day, the samples were soaked in 1:1 sucrose

(30%) and optimal cutting temperature (OCT) compound (Tissue-Tek) solution for 4 h. Samples were then transferred into a cryomold (Tissue-Tek, 25×20×5 mm), covered in OCT and frozen in liquid nitrogen. Using a histology cryotome (Thermo Fisher, CryoStar NX70), sections of 10–40 µm were cut.

*Alizarin Red Staining:* To assess matrix mineralization in cryosections, calcium deposits were stained with Alizarin red. A staining solution (2 mg $ml^{-1}$ Alizarin red S (Sigma-Aldrich, A5533-25G) in distilled water, pH adjusted to 4.12) was applied to the cryosections after two washes with distilled water. The samples were stained for 30 min at room temperature and then washed 5 times until the water came out clear. Imaging at 5× magnification was performed on Leica DMi1 microscope. A color threshold was applied by selecting only the red channel in RGB color space in Fiji/ImageJ. The area of red color was then measured for each condition.

*Osteocalcin Staining:* For osteocalcin immunostaining, cryosections were used. Non-specific antibody binding was blocked with 1% BSA w/v and 5% serum v/v from the host of the secondary antibody for 1 h (goat serum). Primary antibodies were diluted in PBS containing 1% BSA. Immunostaining of osteocalcin was performed using anti-osteocalcin (1:200, Abcam, ab93876) overnight at 4°C. Samples were washed 3×5 min with PBS before incubation with the secondary antibody (1:500 Goat anti-rabbit IgG H&L Alexa 555) for 1 h. Immunohistochemistry staining was validated by a secondary antibody control without adding the primary antibody. Samples were mounted with Mowiol. Sections were imaged using confocal microscopy with a 63× oil immersion objective. To quantify osteocalcin expression, the fluorescence intensity per image was measured as integrated density in Fiji/ImageJ followed by normalization by the cell number in each image.

*Quantification of Permeability:* In order to quantify the permeability of the macroporous PEG hydrogels, the method described by Moreno-Arotzena et al.[2] was adapted. Acellular PEG hydrogels and collagen type I hydrogels were used. 2 mg $ml^{-1}$ collagen type I gel was prepared from an 8.91 mg $ml^{-1}$ stock solution (rat-tail, Corning, 354249) as described by Shin et al.[3] Hydrogels were casted into the channel of a µ-Slide I Luer (Ibidi, channel height: 0.4 mm). To determine the permeability, all hydrogels were first hydrated in PBS for 24 h after crosslinking. Medium reservoirs were filled with 60 µl DMEM and 1 ml syringe barrels without plunger were attached to the luer connectors. 0.5 ml DMEM were added to one of the barrels and 0.1 ml to the other one creating a height and pressure difference that caused interstitial flow through the porous hydrogel. The heights $h_1$ and $h_2$ were measured to calculate the difference in height $h_d$. From this, the pressure difference $\Delta P$ was calculated using **Equation 1**, where $g$ is the gravitational acceleration and $\rho$ is the density of the medium ($\rho_{DMEM}$=1000 kg $m^{-3}$).

$$\Delta P = \rho \times g \times h_d \quad (1)$$

The change of height was measured every 15 min for the first hour and then every 30 min for 4 h in total. An exponential function was fitted to the data according to $\Delta P(t)=P(0)\times e^{-ct}$ to determine the exponent $c$. Since $\Delta P$ changes rapidly within the first hour, only data from time-points 0.75–4 h was considered. Permeability $K$ and the constant $c$ are related according to Darcy's law in **Equation 2**, where $\mu$ is the viscosity of DMEM (7.8×$10^{-4}$ Pa s), $l$ is the length of the hydrogel channel (1.70×$10^{-2}$ m), $A_r$ is the cross section of the syringe barrel and $A$ is the cross section of the hydrogel channel in the direction of fluid flow.

$$K = \frac{c \times \mu \times l \times A_r}{\rho \times g \times A} \quad (2)$$

*Computational Fluid Dynamics (CFD) Simulation:* Confocal images of rhodamine-labeled PEG hydrogel on chip (AIM Biotech, DAX-1) were processed, and the pore geometry was reconstructed using Seg3D (University of Utah, UT, USA). To quantify the FSS within the scaffold that has highly irregular porous geometries, a multiscale and multiphase CFD model previously developed[4] was used. The model involves 2 scales, i.e. (i) the global scale that represents the whole microfluidic chip and (ii) the local scale that models the detailed micro-structures of subsections (dimension: 20×20×30 µm, *n*=4) from the whole scaffold. In the global model, the scaffold region was modelled as porous media with a permeability of $8.67\times10^{-15}$ $m^2$, which was obtained from experimental measurement of a hydrogel with a composition matching the confocal microscopy data. To approximate the experimental condition for dynamic cell culture, two types of flow rates (i.e., 10 µl $min^{-1}$ and 100 µl $min^{-1}$ per port) were applied to the global model as inlet and outlet boundary conditions. Mass flux conservation was applied to the interface between porous media and free fluid. The global model was meshed with 450410 tetrahedral elements. The pressure gradient that was calculated from the global model was applied to the local CFD model for simulating the shear stress on PEG scaffold surfaces. The boundary conditions of local CFD model are shown in Figure 6f. In the local model, the fluid domain of each subsection was meshed by a uniform tetrahedral element size of 0.4 µm, which generated 1322072, 1135226, 1247121, and 1302269 elements, respectively, for 4 discretized subsections. In this study, the fluid was modelled as laminar flow with the dynamic viscosity of DMEM ($7.8\times10^{-4}$ Pa s). The CFD models were solved by a finite volume method (FVM) using ANSYS CFX (ANSYS Inc., PA, USA) under the convergence criteria of root-mean-square residual of the mass and momentum $<10^{-4}$.

### Supplementary Figures

**Supplementary Table 1.** Overview of PEG hydrogel compositions used in this manuscript.

| Figure | 4-PEG-VS [%] | crosslinker | SH/ene | dextran [%] | dextran $M_w$ | HA [%] | RGD | cell density [ml$^{-1}$] |
|---|---|---|---|---|---|---|---|---|
| **2b** | 2.5 (labeled) | PEG-2-SH 2.0 kDa | 0.80 | 1.0 | 40 | 0.50 | GRCGRGDSPG | - |
| **2c** | 2.0–2.5 | KCGPQGIWGQCK, PEG-2-SH 2.0 kDa | 0.80 | 1.0 | 40 | 0.50 | GRCGRGDSPG | - |
| **2d** | 2.0 | GCRDGPQGIWGQDRCG, PEG-2-SH 3.4 kDa | 1.25 (MMP), 0.80 (PEG-2-SH) | 0.0–1.0 | 500 | 0.50 | CGRGDSP | - |
| **2e** | 2.0 | KCGPQGIWGQCK, PEG-2-SH 2.0 kDa | 0.80 | 1.0 | 40 | 0.50 | CGRGDSP | - |
| **3a-c** | 2.0 (labeled) | KCGPQGIWGQCK | 0.80 | 0.0–2.0 | 40 | 0.50 | - | - |
| **3d–f** | 2.0 (labeled) | KCGPQGIWGQCK | 0.80 | 1.0 | 40, 500 | 0.50 | - | - |
| **4b** | 2.0 | GCRDGPQGIWGQDRCG, PEG-2-SH 2.0 kDa | 0.80 | 1.0 | 40 | 0.50 | GRCGRGDSPG | $5.0\times10^5$ |
| **4c** | 2.0 | GCRDGPQGIWGQDRCG, PEG-2-SH 2.0 kDa | 0.80 | 1.0 | 40 | 0.50 | GRCGRGDSPG | $5.0\times10^5$ |
| **4d** | 2.0 | GCRDGPQGIWGQDRCG, PEG-2-SH 2.0 kDa | 0.80 | 1.0 | 40 | 0.50 | GRCGRGDSPG | $3.5\times10^6$ |
| **4e–h** | 2.0 | GCRDGPQGIWGQDRCG, PEG-2-SH 3.4 kDa | 1.25 (MMP), 0.80 (PEG-2-SH) | 1.0 | 500 | 0.50 | CGRGDSP | $3.0\times10^6$ |
| **5a–e** | 2.0 | GCRDGPQGIWGQDRCG, PEG-2-SH 2.0 kDa | 0.80 | 1.0 | 40 | 0.50 | GRCGRGDSPG | $3.5\times10^6$ |
| **6a,b** | 2.2 | KCGPQGIWGQCK | 0.80 | 1.0 | 40 | 0.50 | GRCGRGDSPG | - |
| **6c–e** | 2.0 | GCRDGPQGIWGQDRCG | 1.25 | 1.0 | 40, 500 | 0.50 | CGRGDSP | - |
| **6f–h** | 2.2 | KCGPQGIWGQCK | 0.80 | 1.0 | 40 | 0.50 | CGRGDSP | - |
| **7a** | 2.0 | KCGPQGIWGQCK, PEG-2-SH 3.4 kDa | 0.80 | 1.0 | 500 | 0.50 | CGRGDSP | $3.0\times10^6$ |
| **7b** | 2.2 | KCGPQGIWGQCK | 0.80 | 1.0 | 40 | 0.50 | CGRGDSP | $5.0\times10^5$, $1.0\times10^6$ |
| **7c** | 2.2 | KCGPQGIWGQCK | 0.80 | 1.0 | 40 | 0.50 | CGRGDSP | $1.0\times10^6$ |
| **S1** | 2.0 (labeled) | PEG-2-SH 2.0 kDa | 0.80 | 0, 1 | 40 | 0.50 | - | - |
| **S2a** | 2.0 | PEG-2-SH 2.0 kDa | 0.80 | 1.0 | 40, 500 | 0.50 | - | - |
| **S2b** | 2.5 | PEG-2-SH 2.0 kDa | 0.80 | 1.0 | 40 | 0.25–0.83 | - | - |
| **S3** | 2.0 (labeled) | KCGPQGIWGQCK | 0.80 | 0.0–2.0 | 40 | 0.50 | - | - |
| **S4** | 2.0 (labeled) | KCGPQGIWGQCK | 0.80 | 1.0 | 40, 500 | 0.50 | - | - |
| **S5** | 2.0 (labeled) | KCGPQGIWGQCK | 0.80 | 1.0 | 40, 500 | 0.50 | - | - |
| **S6** | 2.0 | KCGPQGIWGQCK, PEG-2-SH 2.0 kDa | 0.80 | 1.0 | 40 | 0.50 | CGRGDSP | $2.0\times10^6$ |
| **S7** | 2.0 | GCRDGPQGIWGQDRCG, PEG-2-SH 2.0 kDa | 0.80 | 1.0 | 40 | 0.50 | GRCGRGDSPG | $3.5\times10^6$ |
| **S8** | 2.0 | GCRDGPQGIWGQDRCG, PEG-2-SH 3.4 kDa | 1.25 (MMP), 0.80 (PEG-2-SH) | 1.0 | 500 | 0.50 | CGRGDSP | $3.0\times10^6$ |
| **S9** | 2.0 | GCRDGPQGIWGQDRCG, PEG-2-SH 3.4 kDa | 1.25 (MMP), 0.80 (PEG-2-SH) | 1.0 | 500 | 0.50 | CGRGDSP, - | $3.0\times10^6$ |
| **S10** | 2.0 | GCRDGPQGIWGQDRCG | 1.25 | 0.2, 1.0 | 500 | 0.50 | CGRGDSP | $3.0\times10^6$ |
| **S11** | 2.0 | GCRDGPQGIWGQDRCG, PEG-2-SH 2.0 kDa | 0.80 | 1.0 | 40 | 0.50 | GRCGRGDSPG | $3.5\times10^6$ |
| **S12** | 2.2 | KCGPQGIWGQCK | 0.80 | 1.0 | 40 | 0.50 | GRCGRGDSPG | - |

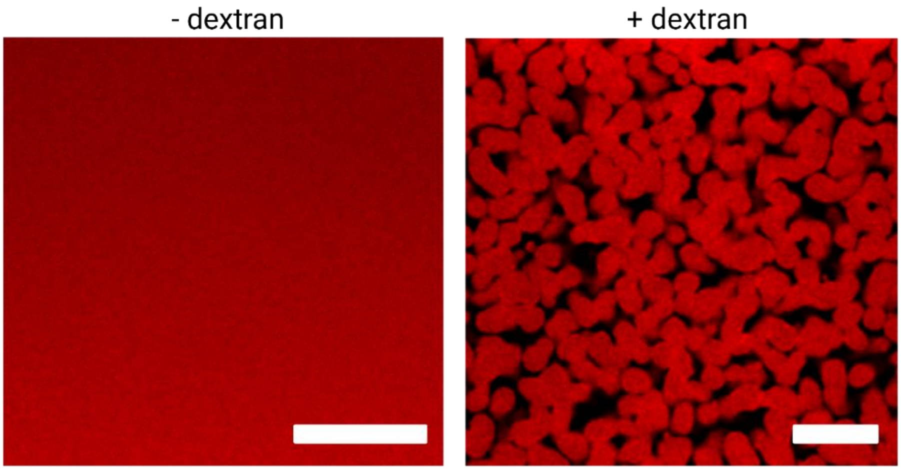


**Supplementary Figure 1.** Confocal microscopy images of PEG hydrogels formed by Michael addition crosslinking without and with dextran, scale bars: 10 µm.

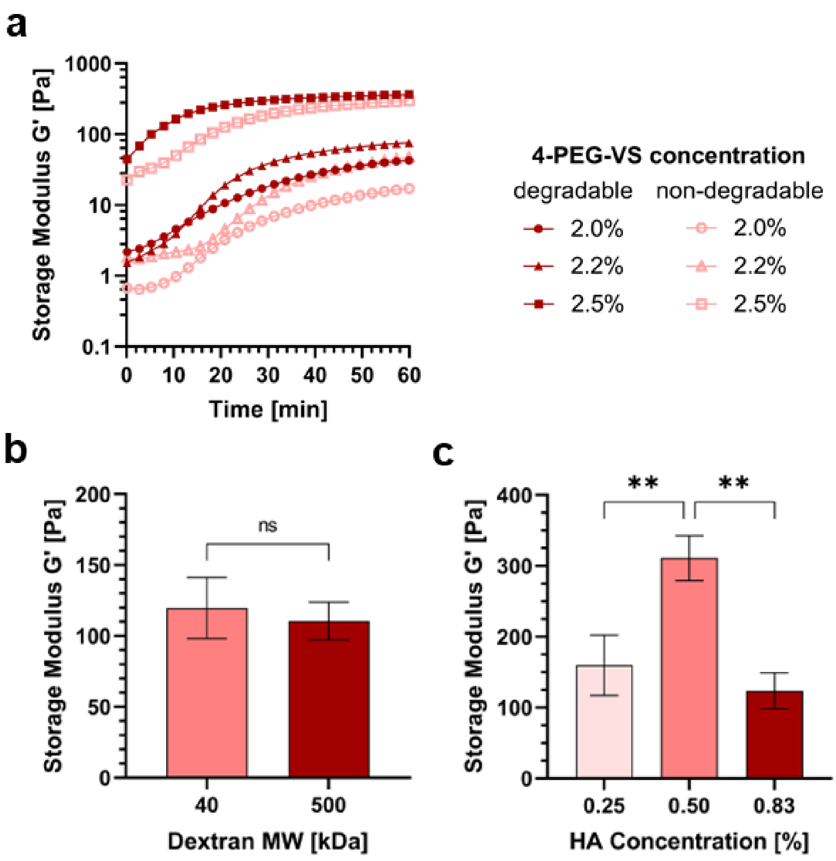


**Supplementary Figure 2.** Rheology of PEG hydrogels. **a)** Representative time-sweep rheology plots of PEG hydrogel compositions with varying PEG concentration during gelation at 37˚C. **b)** Storage modulus (*G′*) of PEG hydrogels with low (40 kDa) and high (500 kDa) dextran $M_W$ after 60 min of crosslinking at 37°C, *n*=3 (Student's t-test). **c)** *G′* of PEG hydrogels with different hyaluronic acid (HA) concentration after 60 min of crosslinking at 37°C, *n*=3 (***p*>0.01, one-way ANOVA/Tukey).

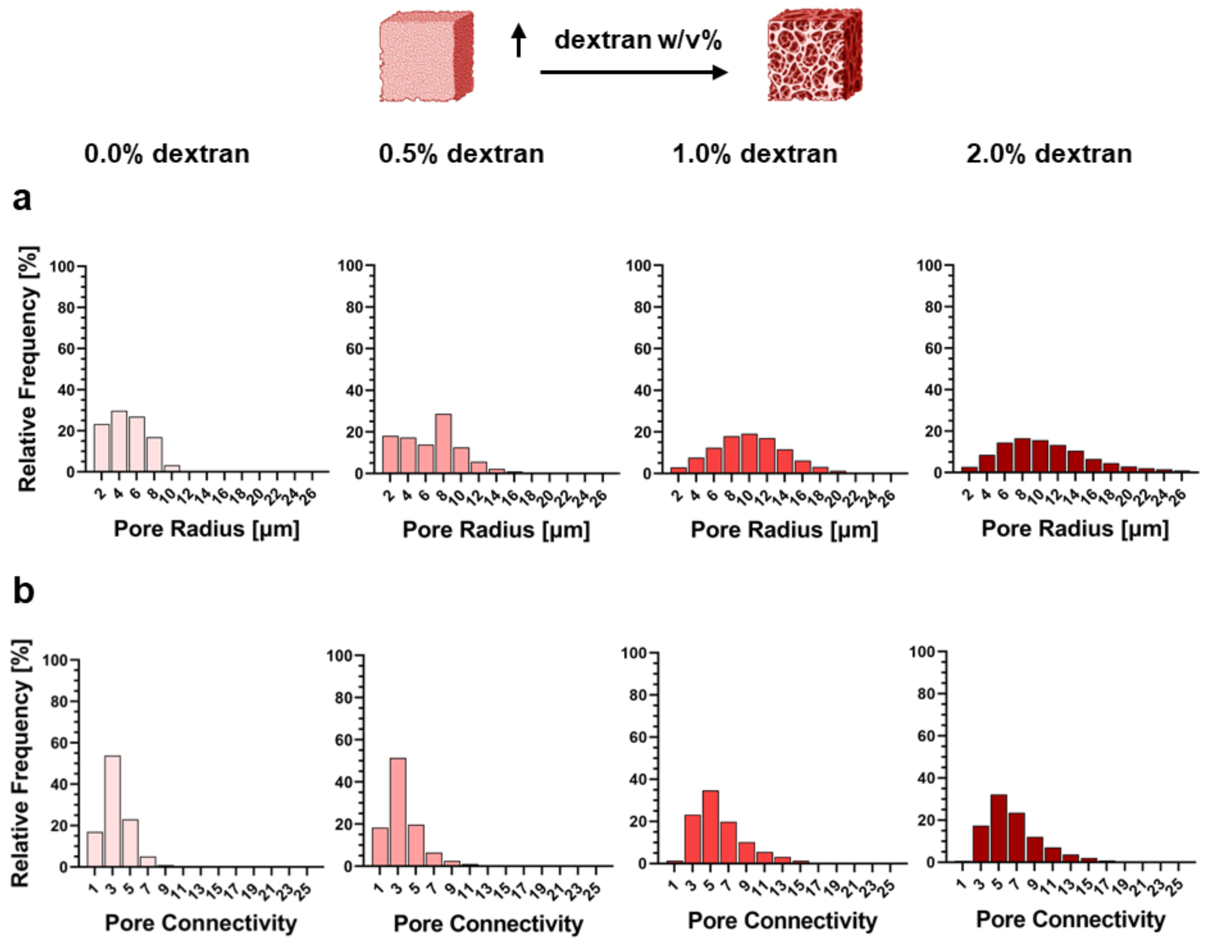


**Supplementary Figure 3.** Characterization of the porous architecture of void-forming PEG hydrogels with varying dextran concentration. **a)** Distribution of pore radii in PEG hydrogels with dextran concentrations ranging from 0.0–2.0%, *n*=3. **b)** Distribution of pore connectivity in PEG hydrogels with dextran concentrations ranging from 0.0–2.0%, *n*=3.

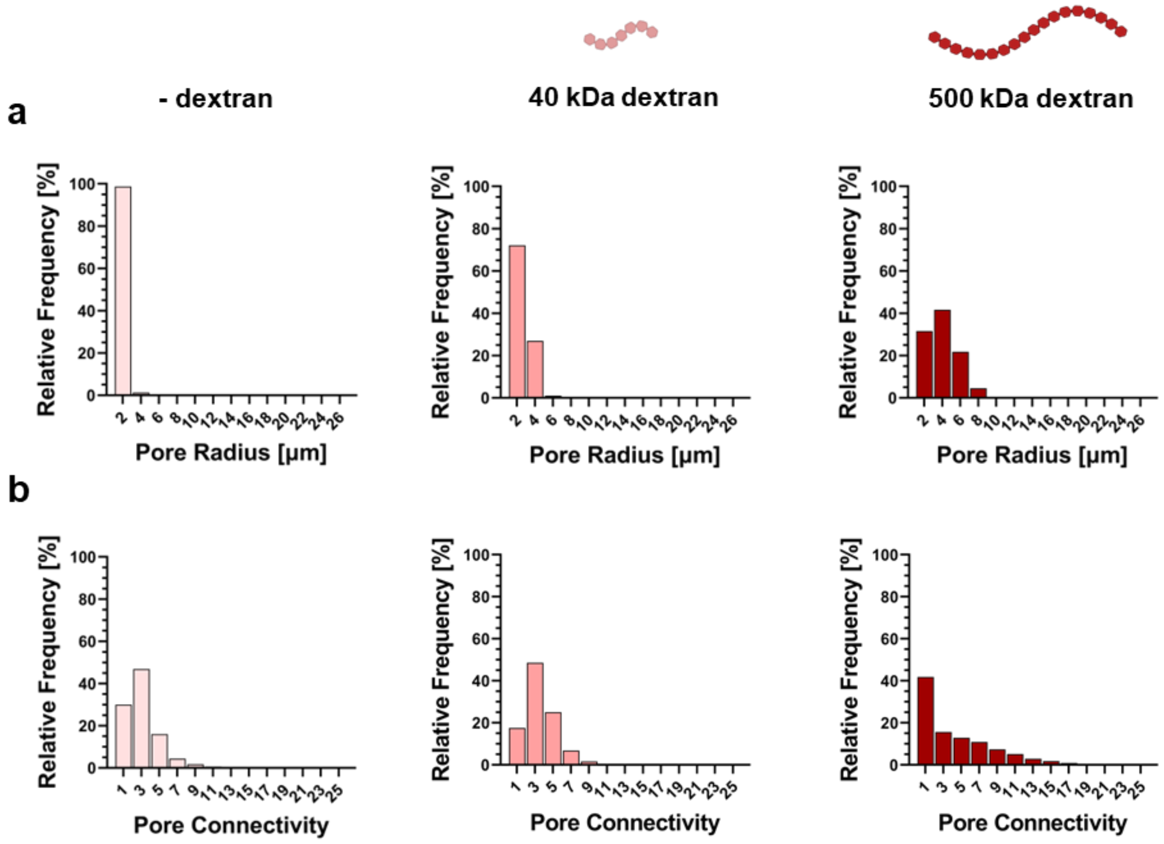


**Supplementary Figure 4.** Characterization of the porous architecture of void-forming PEG hydrogels with low (40 kDa) and high (500 kDa) dextran $M_W$ or without the addition of dextran. **a)** Distribution of pore radii in PEG hydrogels as a function of hydrogel composition, *n*=3. **b)** Distribution of pore connectivity in PEG hydrogels as a function of hydrogel composition, *n*=3.

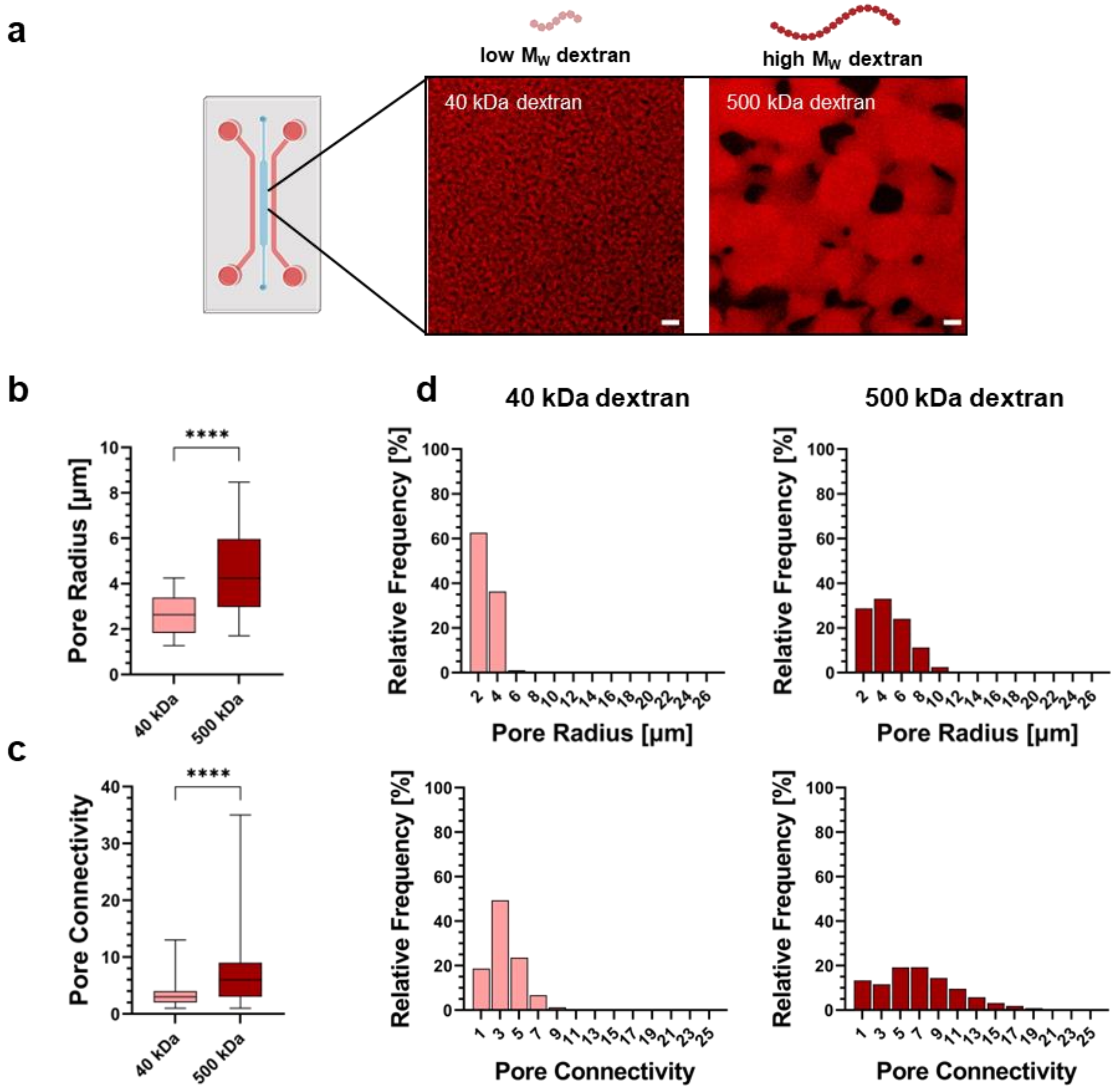


**Supplementary Figure 5.** Characterization of the porous architecture of void-forming PEG hydrogels on a microfluidic chip. **a)** Confocal microscopy images of rhodamine-labeled PEG hydrogels formed with 1.0% low $M_W$ (40 kDa) and high $M_W$ (500 kDa) dextran, scale bars: 10 µm. **b-c)** Quantification of pore radius and pore connectivity of hydrogels formed with low $M_W$ (40 kDa) and high $M_W$ (500 kDa) dextran on chip, $n$=3 (****$p$>0.0001, Student's t-test). **d)** Distribution of pore radii (top) and pore connectivity (bottom) in PEG hydrogels with low and high $M_W$ dextran on a microfluidic chip, $n$=3.

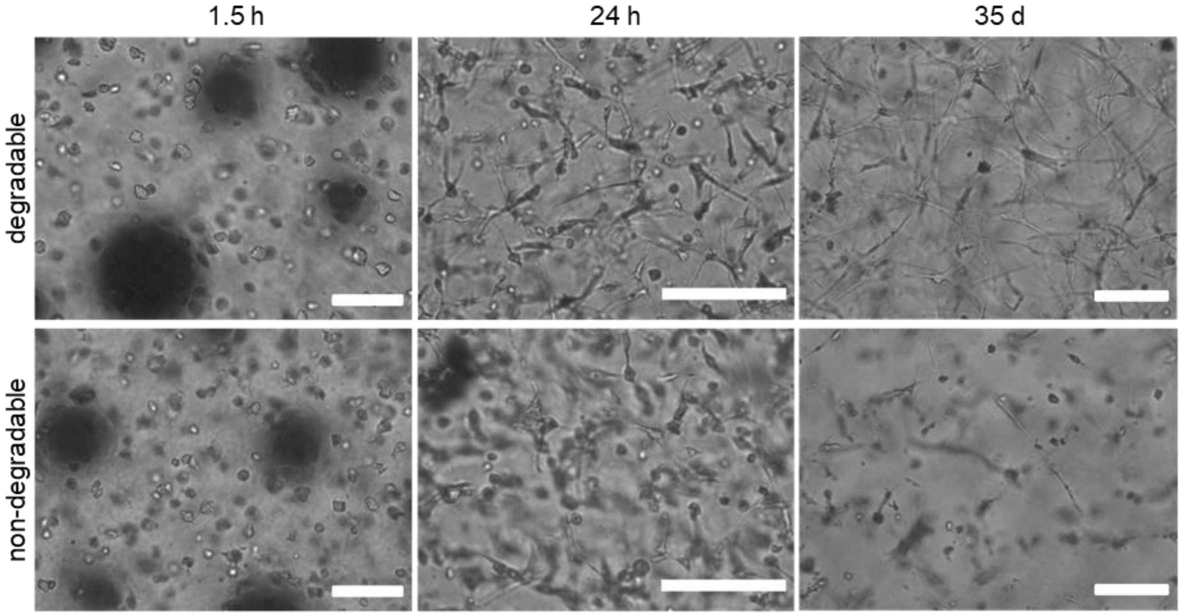


**Supplementary Figure 6.** Microscopy images of cell morphologies within degradable and non-degradable PEG hydrogels showing human mesenchymal stem cell (hMSC) spreading after 1.5 h, network formation after 1 day. The cell network was stable in degradable hydrogels, whereas network degeneration was observed in non-degradable hydrogels following osteogenic cultivation for 35 days, scale bars: 50 µm.

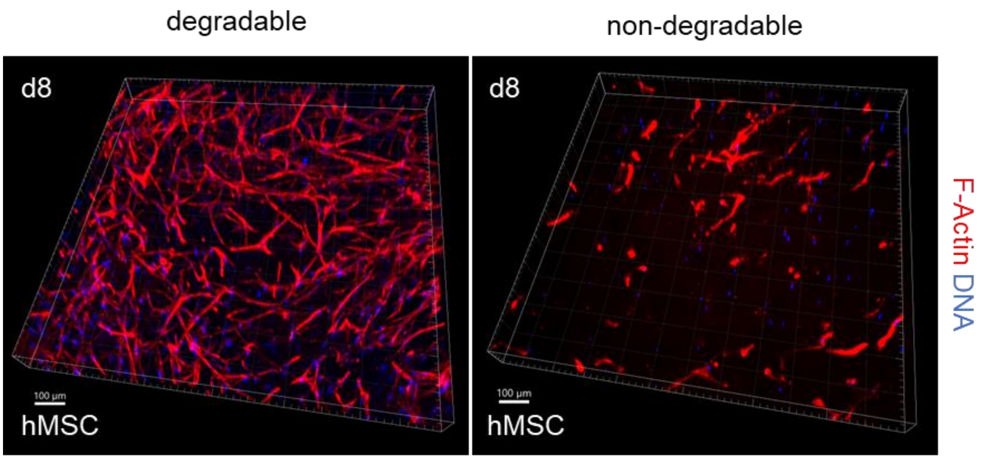


**Supplementary Figure 7.** Representative 3D view of actin-nuclei-stained hMSC networks in degradable and non-degradable hydrogels after 8 days of culture, scale bars: 100 µm.

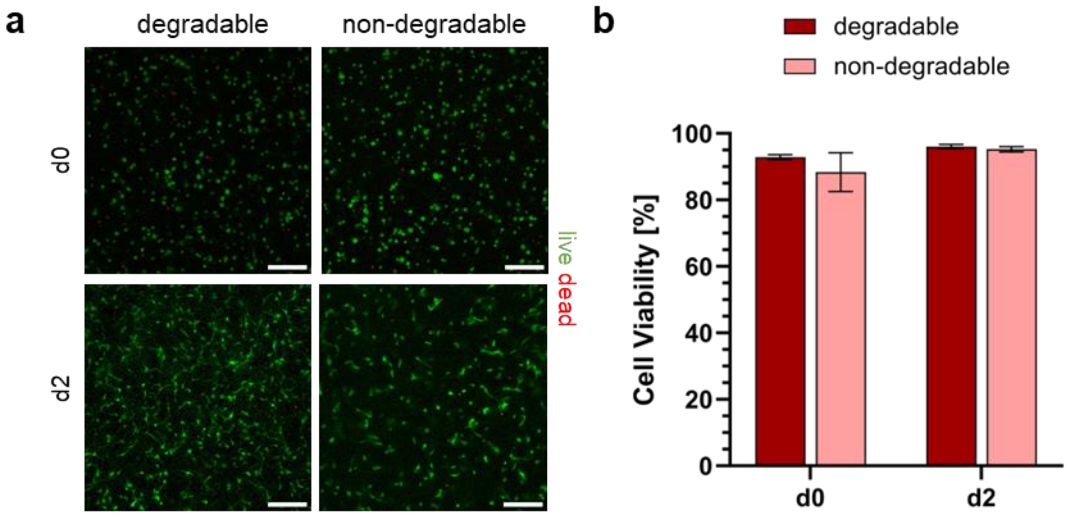


**Supplementary Figure 8.** Cell viability of human osteoblasts (hOBs) after embedding and 2 days of osteogenic culture within degradable and non-degradable PEG hydrogels. **a)** Confocal microscopy images (MIPs) of live/dead staining, scale bars: 100 µm. **b)** Quantification of cell viability based on live/dead staining, *n*=3 (two-way ANOVA/Tukey).

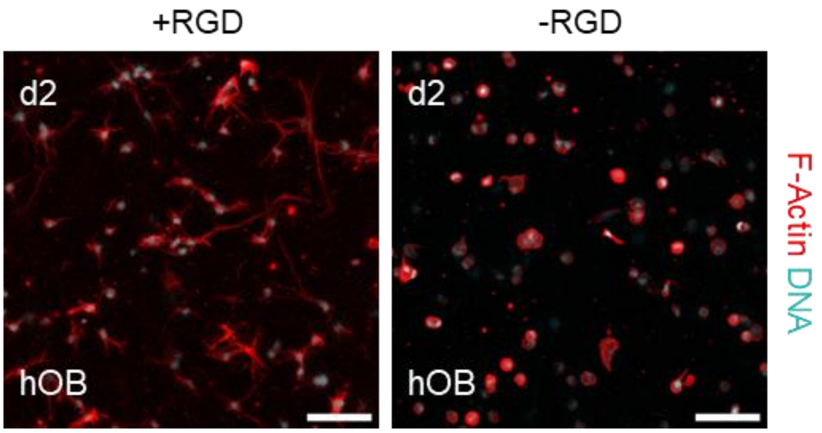


**Supplementary Figure 9.** Confocal microscopy images of actin-nuclei-stained hOBs following 2 days of osteogenic cultivation in matrix metalloproteinase (MMP)-degradable PEG hydrogels, emphasizing the importance of RGD motifs for 3D cell network formation, scale bars: 100 µm.

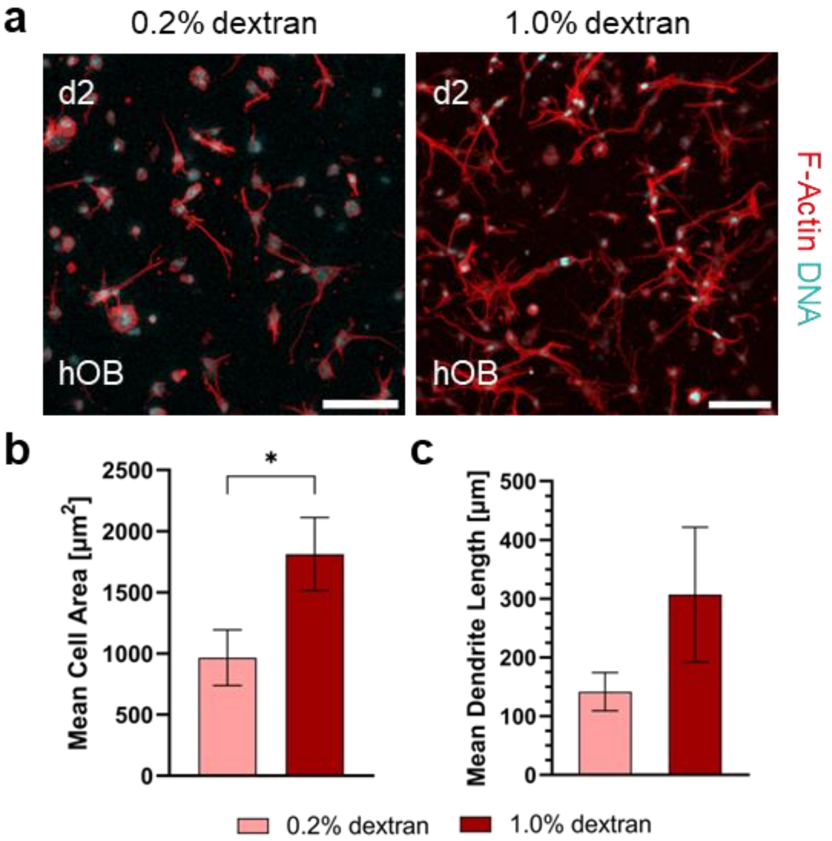


**Supplementary Figure 10. a)** Confocal microscopy images of actin-nuclei-stained hOBs following 2 days of osteogenic cultivation in MMP-degradable PEG hydrogels, emphasizing the importance of dextran concentration and resulting pore size for 3D cellular network formation, scale bars: 100 µm. **b)** Quantification of mean cell area in hydrogels with 0.2% and 1.0% dextran, *n*=3 (**p*<0.05, Student's t-test). **c)** Quantification of mean dendrite length per cell using NeuriteQuant in hydrogels with 0.2% and 1.0% dextran, *n*=3 (Student's t-test).

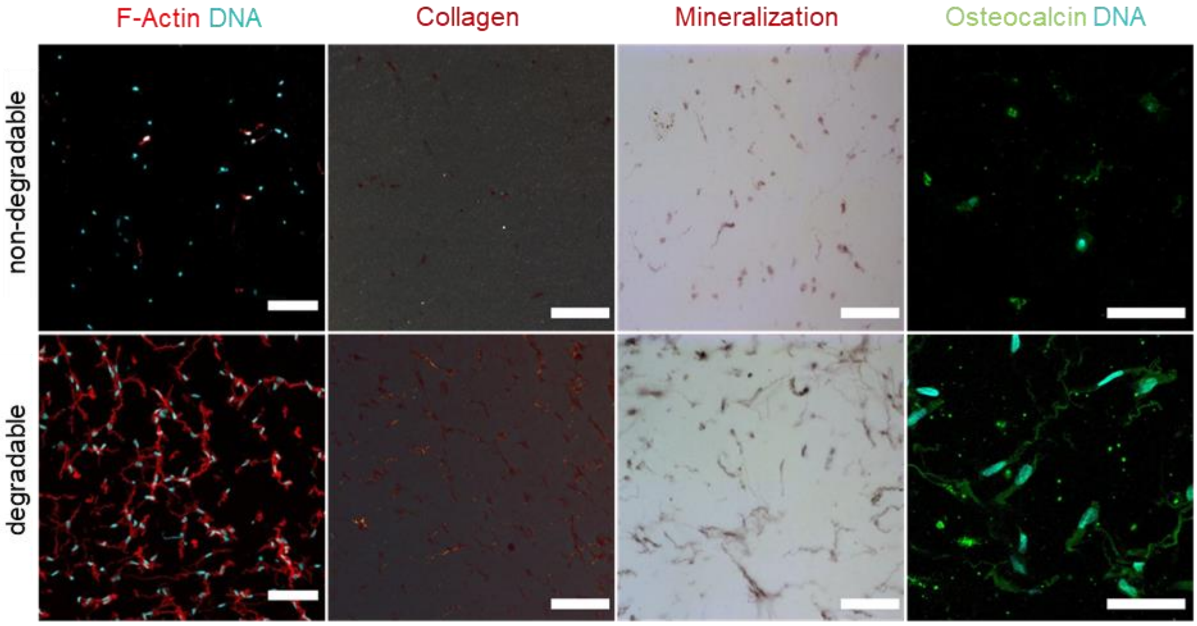


**Supplementary Figure 11.** Histological analysis of static hMSC culture within MMP-degradable and non-degradable PEG hydrogels on day 8. Microscopy images of osteogenic markers, including cell morphology determined by confocal microscopy (MIP, scale bars: 100 µm), collagen fiber secretion determined by Picrosirius-polarization microscopy (scale bars: 100 µm), matrix mineralization determined by Alizarin red staining (scale bars: 100 µm) and osteocalcin expression by immunohistostaining (MIP, scale bars: 50 µm).

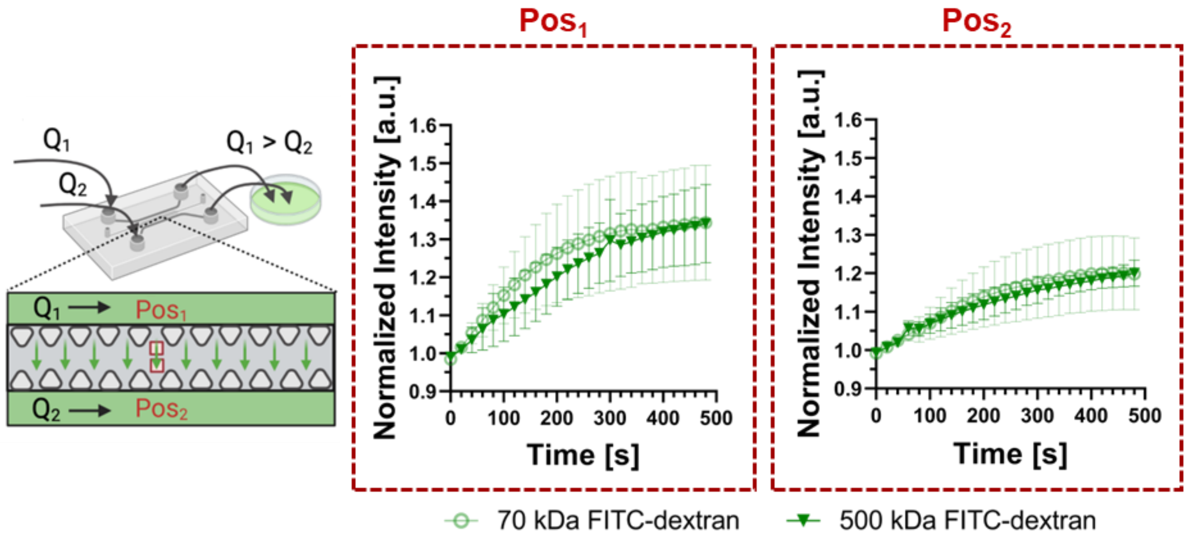


**Supplementary Figure 12.** Microfluidic setup to visualize fluid flow through PEG hydrogels, $Q_1$ and $Q_2$ denote volumetric flow rate, $Pos_1$ and $Pos_2$ represent positions of image acquisition (left). Changes in normalized fluorescent intensity over time in positions $Pos_1$ and $Pos_2$, $n$=3 (right).

### Supplementary Movies

**Supplementary Movie 1.**

**Pore formation by PIPS in PEG hydrogel.** Time-lapsed confocal microscopy showing *in situ* formation of macroporous PEG hydrogels by phase separation between labeled 4-PEG-VS and dextran during crosslinking at 37°C, scale bar: 10 µm.

**Supplementary Movie 2.**

**Macroporous PEG hydrogel after crosslinking.** Confocal microscopy images of macroporous PEG hydrogels after crosslinking and washing in PBS, scale bar: 10 µm.

**Supplementary Movie 3.**

**hMSC network within MMP-degradable PEG hydrogel.** 3D animation of an hMSC-derived cell network within a macroporous MMP-degradable PEG hydrogel on day 8: F-actin (red) and nuclei (blue). The video was created with IMARIS.

**Supplementary Movie 4.**

**Visualization of fluid flow through macroporous PEG hydrogel on a microfluidic chip.** Time-lapsed fluorescence microscopy images showing fluorescein isothiocyanate (FITC)-dextran tracer molecules (500 kDa) perfusing through a macroporous MMP-degradable PEG hydrogel on-chip in response to a pressure gradient, scale bar: 200 µm.
